# Mapping transcription factor occupancy using minimal numbers of cells *in vitro* and *in vivo*

**DOI:** 10.1101/158931

**Authors:** Luca Tosti, James Ashmore, Boon Siang Nicholas Tan, Benedetta Carbone, Tapan K Mistri, Valerie Wilson, Simon R. Tomlinson, Keisuke Kaji

## Abstract

The identification of transcription factor (TF) binding sites in the genome is critical to understanding gene regulatory networks (GRNs). While ChIP-seq is commonly used to identify TF targets, it requires specific ChIP-grade antibodies and high cell numbers, often limiting its applicability. <u>D</u>NA <u>a</u>denine <u>m</u>ethyltransferase <u>id</u>entification (DamID), developed and widely used in *Drosophila*, is a distinct technology to investigate protein-DNA interactions. Unlike ChIP-seq, it does not require antibodies, precipitation steps or chemical protein-DNA crosslinking, but to date it has been seldom used in mammalian cells due to technical impediments. Here we describe an optimised DamID method coupled with next generation sequencing (DamID-seq) in mouse cells, and demonstrate the identification of the binding sites of two TFs, OCT4 and SOX2, in as few as 1,000 embryonic stem cells (ESCs) and neural stem cells (NSCs), respectively. Furthermore, we have applied this technique *in vivo* for the first time in mammals. Oct4 DamID-seq in the gastrulating mouse embryo at 7.5 days post coitum (dpc) successfully identified multiple Oct4 binding sites proximal to genes involved in embryo development, neural tube formation, mesoderm-cardiac tissue development, consistent with the pivotal role of this TF in post-implantation embryo. This technology paves the way to unprecedented investigations of TF-DNA interactions and GRNs in specific cell types with limited availability in mammals including *in vivo* samples.

## Introduction

Genome-wide transcription factor (TF) occupancy is commonly assessed by chromatin immunoprecipitation followed by next-generation sequencing (ChIP-seq), but the need for both ChIP-grade antibodies and high numbers of cells limits its utility (Barski et al. 2007). Although recent improvements of the ChIP-seq protocol, such as the iChIP and the ChIPmentation strategies (Lara-Astiaso et al. 2014; Schmidl et al. 2015), enabled the detection of histone marks across the genome using only 500 cells, they still require 10,000-500,000 cells to identify targets of the TF PU.1. The use of CETCh-seq (CRISPR epitope tagging followed by ChIP-seq) overcomes the need for TF-specific ChIP-grade antibodies, but it does not reduce the required numbers of cells (Savic et al. 2015). <u>D</u>NA <u>a</u>denine <u>m</u>ethyltransferase Identification (DamID) is a distinct method to investigate protein-DNA interactions, based on the exogenous expression of a protein of interest (POI) tethered to the DNA adenine methyltransferase (Dam) from *E. Coli* (van Steensel et al. 2001). Dam specifically methylates the adenine within a GATC sequence; hence the expression of the Dam-POI fusion protein creates unique GA^me^TC marks on DNA adjacent to the POI binding sites. Subsequent genomic DNA extraction, digestion with the GA^me^TC-specific restriction enzyme DpnI, adapter ligation, PCR amplification and microarray (DamID-chip) or next generation sequencing (DamID-seq) allow identification of the POI binding events. Unlike ChIP-seq, this technique does not require formaldehyde fixation or immunoprecipitation steps that could lead to data biases (Baranello et al. 2016) or loss of materials. DamID has been used in seminal studies in *Drosophila* for over 100 chromatin proteins and transcription factors (van Bemmel et al. 2013; Moorman et al. 2006; Filion et al. 2010). However, only limited success has been reported in mammalian cells probably due to technical impediments. In particular, very low expression of the Dam protein without tethering POI (Dam-only) is sufficient to methylate DNA (Wines et al. 1996) since Dam itself can bind DNA and has highly processive methylation activity (Urig et al. 2002). The detection of POI-specific binding sites in DamID depends on the comparison of methylation signatures between Dam-only and Dam-POI expressing cells. Thus, expressing Dam-only and Dam-POI at equally low level in two independent populations is critical to identify POI-dependent methylation signal. This issue becomes even more relevant when the POI interacts with DNA at open chromatin loci (such as TFs), since Dam-only also preferentially binds and methylates nucleosome free DNA (van Steensel et al. 2001). Since the first mammalian HP1β DamID-chip paper (Vogel et al. 2006), only a handful of publications have reported the use of DamID-chip/seq for TFs in mammalian cells (Supplemental Table S1). We have overcome the above mentioned challenges by applying translation reinitiation-mediated DamID, recently reported in *Drosophila* (Southall et al. 2013), to a mouse system. In combination with Tn5 transposase-mediated tagmentation and next generation sequencing, this novel DamID-seq enabled us to detect clear TF binding signatures with as little as 1,000 cells. This work details the improvements of the DamID-seq technology and demonstrates for the first time the identification of *in vivo* OCT4 binding sites in the gastrulation stage mouse embryo.

## Results

### Development of a translation reinitiation-mediated DamID-seq in mouse cells

In the original protocol for mammalian DamID-chip, Dam-only and Dam-HP1β were expressed *via* plasmid transfection under the ecdysone-inducible promoter (Vogel et al. 2006). The leakiness of this promoter (i.e. in the absence of ecdysone) was sufficient to achieve an optimal, low expression of Dam-only/POI and this strategy has been used for many DamID experiments in *Drosophila* Kc cells (Filion et al. 2010; van Bemmel et al. 2013). However, this approach limits the applicability of DamID where efficient transfection/viral infection or propagation of transfected cells is possible. In addition, expression levels of Dam-only and Dam-POI depend on transfection/infection efficiency, integration copy numbers and integration sites, hence achieving an equal expression level in two independent samples is technically challenging. Recently the phenomenon of translation reinitiation has been exploited in *Drosophila* to achieve an optimal Dam expression level in a tissue specific manner in combination with the GAL4-UAS system (Southall et al. 2013). Translation reinitiation takes place since eukaryotic ribosome does not always detach from mRNA at the stop codon of a 1^st^ open reading frame (ORF) and can restart translation of a 2^nd^ downstream ORF. Expression levels of the protein encoded by the 2^nd^ ORF decrease as the length of the 1^st^ ORF increases (Kozak 2001), providing a method by which to fine tune the level of Dam-only/Dam-POI expression (Southall et al. 2013).

To optimise translation reinitiation-mediated DamID in mammalian systems, we initially focused on the binding of the master regulator of pluripotency, Oct4, in mouse embryonic stem cells (ESCs). Preceding DamID experiments, the functionality of the Dam-Oct4 fusion protein was confirmed by maintaining an undifferentiated state in an inducible *Oct4* knockout ESC line (Supplemental Fig. S1).

**Supplemental Figure 1:**
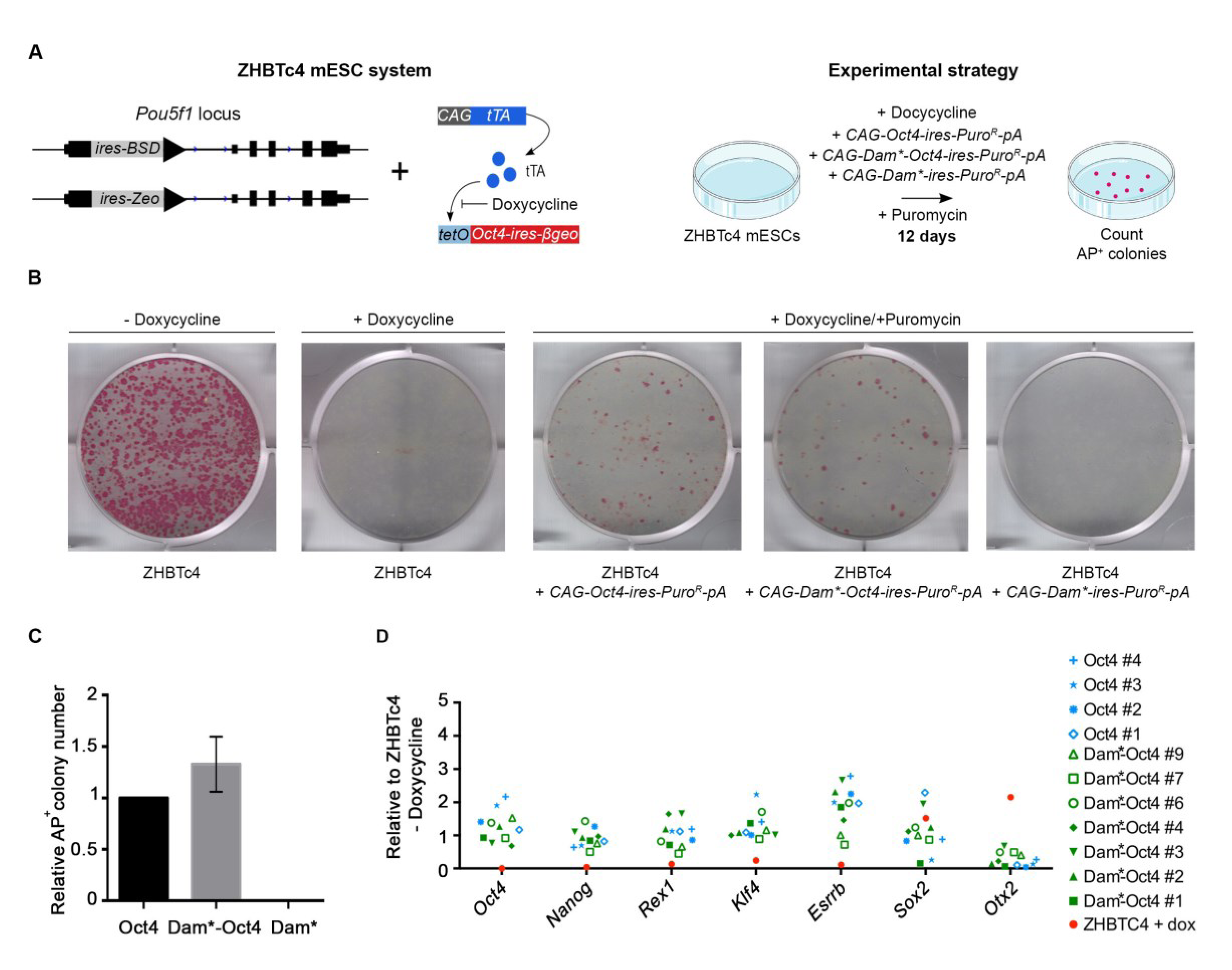
Dam-Oct4 is functionally equivalent to Oct4. (*A*) In ZHBTc4 ESCs, two endogenous *Oct4* alleles were knocked out, and tetO promoter-driven *Oct4* sustains an undifferentiated state in the absence of doxycycline (Niwa et al. 2000) (left). The ability to maintain the undifferentiated state of *Oct4, Dam*-Oct4* and *Dam\** expression vector was assessed upon administration of doxycycline (right). *Dam\** (Dam D181A mutant), which lacks methylation activity without altering its DNA binding capacity (Liebert et al. 2004), was used in these experiments as over-expression of wild-type Dam is toxic. (*B*) Alkaline phosphatase (AP) staining identified undifferentiated ZHBTc4 ESC colonies maintained by wild type *Oct4* and *Dam*-Oct4.* (*C*) Quantification of AP+ colonies in ZHBTc4 ESCs transfected with Oct4 wild type, Dam*-Oct4 or Dam*-only. Data represent mean ± s.e.m. (n=2). (*D*) Gene expression of ZHBTc4 ESC clones (passage 5) maintained with Oct4 wild type (blue), Dam*-Oct4 (green), and differentiated cells without a rescue plasmid (red).

We then generated an ESC line containing a PhiC31 integrase-mediated cassette exchange platform within the *Rosa26* locus (Fig. 1A), allowing us to generate various cell lines with Dam-only/POI expression under the endogenous *Rosa26* promoter with high (~100%) efficiency via simple plasmid transfection and drug selection.

**Figure 1.**
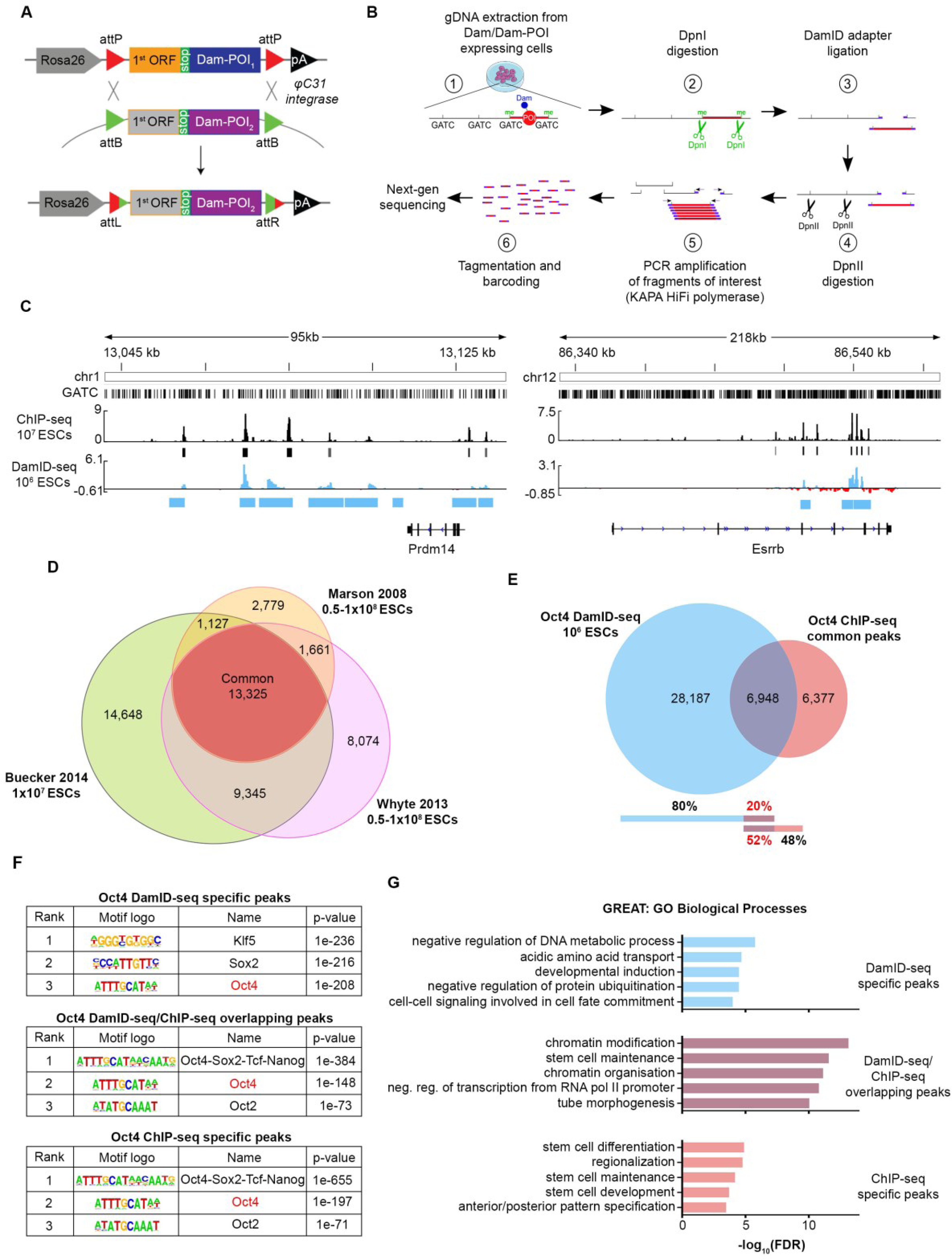
Optimisation of DamID-seq in mouse ESCs and comparison with ChIP-seq. (*A*)φC31 integrase-mediated cassette exchange system used to generate the cell lines in this study. POI = protein of interest; ORF = open reading frame. (*B*) The optimized DamID-seq workflow. (*C*) Oct4 DamID-seq tracks generated from 10^6^ ESCs and Oct4 ChIP-seq tracks generated from 10^7^ ESCs (Buecker et al. 2014). The bars below each track represent ChIP-seq (black) and DamID-seq (blue) statistically significant peaks. Y axis represents read counts per million of Dam-Oct4 (Dam-subtracted) and Oct4 ChIP-seq (input-subtracted), respectively. (*D*) Peak overlap between three different published Oct4 ChIP-seq datasets (Marson et al. 2008; Whyte et al. 2013; Buecker et al. 2014). (*E*) Overlap between Oct4 DamID-seq peaks and ChIP-seq common peaks. (*F*) Motif enrichment analysis (Heinz et al. 2010) of the Oct4-bound peaks identified only by DamID-seq, ChIP-seq or by both technologies. (*G*) Gene ontology enrichment analysis of Oct4 peaks identified only by DamID-seq, ChIP-seq or by both using GREAT (McLean et al. 2010).

To identify the optimal length of the 1^st^ ORF for translation reinitiation-mediated DamID, we placed the Dam-only/Dam-Oct4 coding sequences downstream of the stop codon of three different ORFs: Hygromycin *(Hyg,* 1,032 bp), Neomycin *(Neo,* 804 bp), and Blasticidin *(Bsd,* 393 bp) resistance cassettes (Supplemental Fig. S2). Then we measured the methylation level at the *Pou5f1/Oct4* locus in each cell line by quantitative PCR-based DamID (qDamID) (Supplemental Fig. S2 and Methods).

**Supplemental Figure 2:**
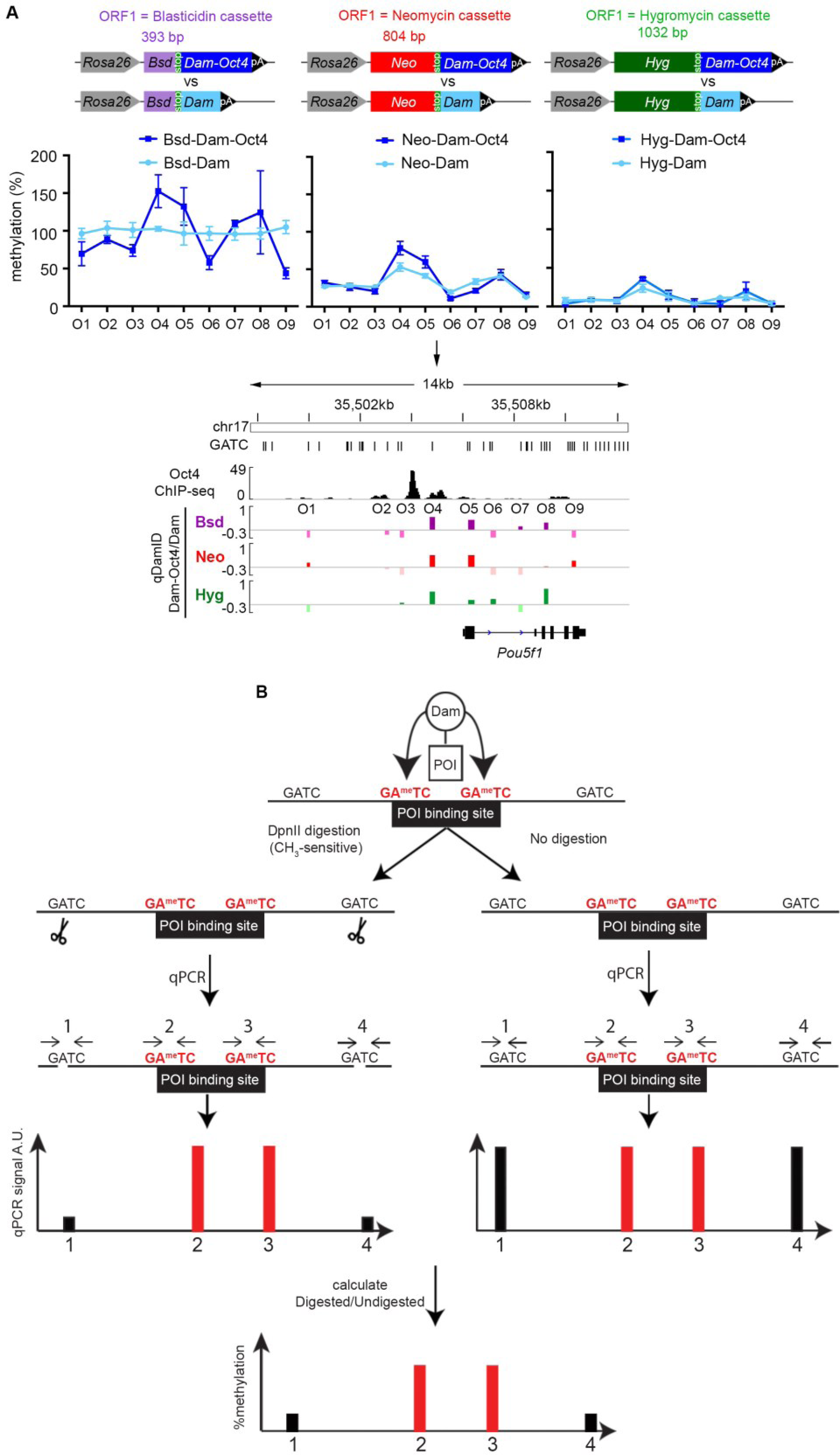
Identification of the optimal 1st ORF for translation reinitiation-mediated DamID. (*A*) Methylation levels in the *Pou5f1* locus measured by qDamID (see *B)* in cell lines with different 1^st^ ORFs before the Dam/Dam-Oct4 coding sequences. Data represent mean ± s.e.m. (n=2). O1-O9 represents the position of the GATCs analysed by qDamID. The methylation levels (%) in *Dam-Oct4*, *Dam* expressing cell lines were used to calculate log_2_(Dam-Oct4/Dam). Bsd = blasticidin resistance gene; Neo = neomycin resistance gene; Hyg = hygromycin resistance gene. (*B*) Schematic representation of the qDamID workflow. POI = protein of interest; GA^me^TC = adenine-methylated GATC sequence; A.U. = arbitrary unit.

We detected positive methylation ratios (Dam-Oct4/Dam-only, i.e. Oct4 binding sites) at known Oct4 binding sites in agreement with ChIP-seq data using the three different cassettes; however, using *Neo* as the 1^st^ ORF produced showed the widest dynamic range of methylation levels without saturation (Supplemental Fig. S2A). Therefore, we used *Neo* as the 1^st^ ORF cassette in subsequent experiments.

### Translation reinitiation-mediated DamID enables the detection of TF binding dynamics during ESC differentiation

One of the big advantages of generating Dam-only/POI expressing ESC lines is that they can be differentiated into any cell type. In fact, the Rosa-Neo-Dam and Rosa-Neo-Dam-Oct4 ESC lines similarly well differentiated into epiblast-like cells (EpiLCs, Supplemental Fig. S3A) and fibroblast-like cells *in vitro.* Since the expression level of Dam-only/POI *via* translation reinitiation is extremely low, it is unlikely to affect the function of the cells.

**Supplemental Figure 3:**
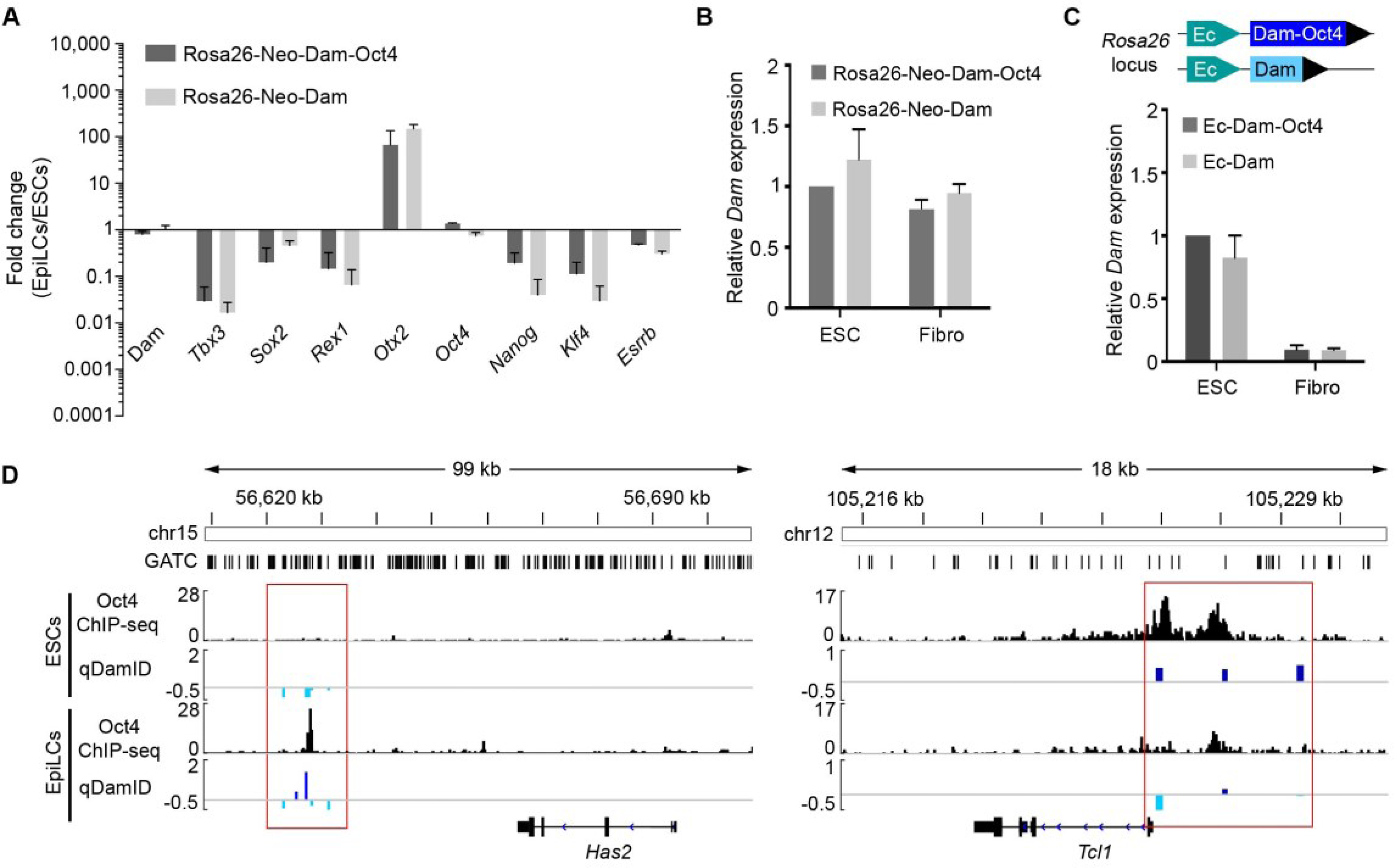
Gain and loss of qDamID signal upon ESCs differentiation. (*A*) Similar expression changes of pluripotency-related genes after conversion of Rosa26-Neo-Dam-Oct4 and Rosa26-Neo-Dam ESCs to EpiLCs. Data plotted as mean ± s.e.m. (n=3). (*B*) Expression level of *Dam* in Rosa26-Neo-Dam-Oct4 and Rosa26-Neo-Dam ESCs and *in vitro* differentiated fibroblasts. Data represent mean ± s.e.m. (n=2). (*C*) Expression level of *Dam* in Rosa26-Ec-Dam-Oct4 and Rosa26-Ec-Dam ESCs and *in vitro* differentiated fibroblasts. Data represent mean ± s.e.m. (n=2). Ec = Ecdysone-inducible promoter. (*D*) qDamID analysis in the *Has2* and *Tcl1* loci with Rosa26-Neo-Dam-Oct4 and Rosa-26-Neo-Dam ESCs and EpiLCs in comparison with Oct4 ChIP-seq data (Buecker et al. 2014).

*Neo-Dam* and *Neo-Dam-Oct4* mRNA levels remained constant after differentiation as we used the ubiquitously active endogenous *Rosa26* promoter (Supplemental Fig. S3B). In contrast, when we placed the ecdysone-inducible promoter-driven *Dam-only/Oct4* cassettes in the *Rosa26* locus, the transgenes were silenced following differentiation into fibroblast-like cells (Supplemental Fig. S3C). Although the Rosa-Neo-Dam and Rosa-Neo-Dam-Oct4 lines constitutively express Dam-only/Dam-Oct4, we could observe loss and gain of the Oct4 binding signatures using a 72-hour EpiLC differentiation protocol (Supplemental Fig. S3D), in agreement with ChIP-seq data (Buecker et al. 2014). These results indicated that the translation re-initiation-based DamID system could detect distinct POI binding signal in different cell types, as long as cells divide and dilute the methylation signal generated in previous differentiation stages.

### Optimization of DamID-seq with 10^6^ ESCs

A few studies have described the use of DamID-seq for the analysis of TF binding in mammalian cells (Supplemental Table S1), yet with little modifications of the original DamID-chip protocol (Vogel et al. 2007). Thus we revisited each step of the protocol and optimized it for DamID-seq (Fig. 1B). Genomic DNA (gDNA) was extracted using the Quick-gDNA™ MicroPrep kit (Zymo Research), and then digested with DpnI. After adapter ligation, DNA was digested with DpnII before adapter-mediated PCR amplification. The DpnII digestion step was described in the original protocol, but it has been omitted in recent studies (Kind et al. 2015; Jacinto et al. 2015; Bouveret et al. 2015). We found that this step was critical to achieve high signal-to-noise ratio in TF DamID-seq. When the DpnII digestion was omitted, the intensity of the POI signal over the Dam-only control was reduced, making the identification of TF binding sites difficult (Supplemental Fig. S4A).

**Supplemental Figure 4:**
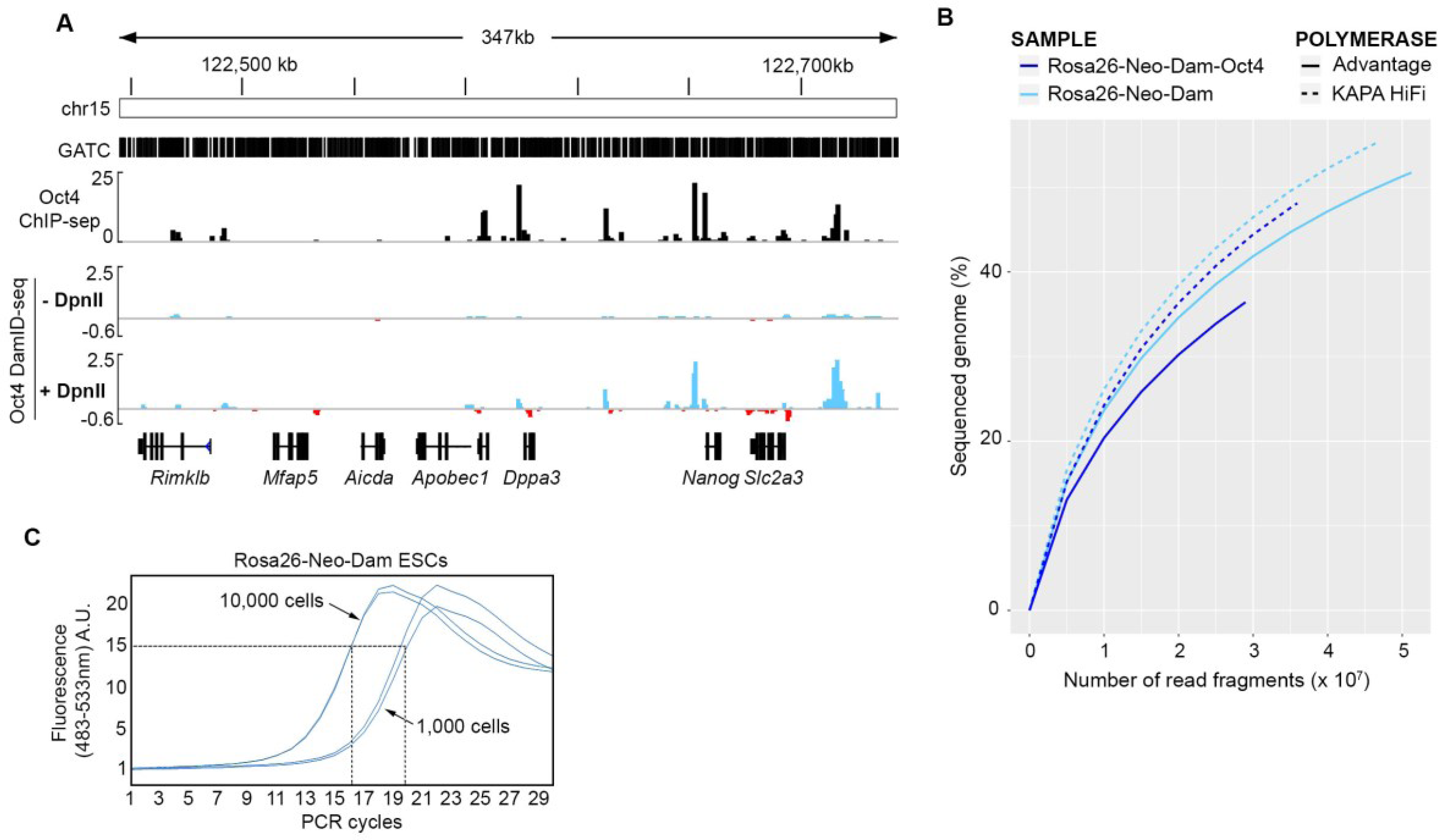
Optimization of the DamID-seq protocol. (*A*) A screenshot of Oct4 DamID-seq tracks generated from 10^4^ ESCs with or without the DpnII digestion step (step #4 in Fig. 1B) in comparison with Oct4 ChIP-seq (Buecker et al. 2014). (*B*) The percentage of sequenced genome when using Advantage2® Polymerase or KAPA HiFi Polymerase in the PCR amplification step (step #5 in Fig. 1B). (*C*) qPCR analysis to estimate the optimal number of PCR cycles required to amplify each sample. Using this approach, it is possible to determine the number of PCR cycles necessary in order to stop the amplification process in the linear phase (e.g. 16 cycles for 10,000 and 20 cycles for 1,000 *Rosa26-Neo-Dam* ESCs).

For the amplification of the adapter ligated DNA fragments, we found that the KAPA HiFi polymerase provided a better genome coverage than Advantage2 polymerase previously used (Southall et al. 2013; Guelen et al. 2008; van Steensel et al. 2001) (Supplemental Fig. S4B). We also introduced a quantitative PCR step to determine the optimal number of PCR cycles for the fragment amplification in order to minimize amplification biases (Supplemental Fig. S4C). All these steps could be performed in a single tube and the amplified DNA was then purified using SPRI magnetic beads. The purified Dam-only/POI target DNA was subjected to library preparation for Illumina sequencing with Tn5 transposition (Picelli et al. 2014), which allowed us to fragment the DNA to the desired size range (~250-350 bp) for NGS and to introduce Illumina sequencing-compatible ends in a 5 minute reaction. This DamID-seq protocol (from gDNA extraction to NGS library preparation) can be accomplished in ~3 days.

We initially performed Oct4 DamID-seq using 10^6^ ESCs. Visual inspection of the Oct4 DamID-seq tracks, generated by subtraction of the Dam-only signal from the Dam-Oct4 signal, revealed good agreement with Oct4 ChIP-seq data generated with 10^7^ ESCs (Fig. 1C). We re-analysed three previously published Oct4 ChIP-seq datasets (Marson et al. 2008; Whyte et al. 2013; Buecker et al. 2014) (Fig. 1D), and used the common 13,325 peaks for further comparison with the 35,135 Oct4 DamID-seq peaks identified in this study using a new DamID-seq data analysis pipeline described in the Methods (the source code is available in the Supplemental Material) (Fig. 1E). About 50% of the Oct4 ChIP-seq peaks overlapped with the DamID-seq peaks, and motif enrichment analysis (Heinz et al. 2010) confirmed that the Oct4 motif was enriched in the overlapping peaks, as well as in DamID-specific and in ChIP-specific peaks (Fig. 1F). Interestingly, DamID-specific peaks were highly enriched with Sox2 and Klf5 motifs, instead of the Oct4-Sox2-Tcf-Nanog binding motif enriched in the overlapping and ChIP-seq specific peaks (Fig. 1F). Oct4 and Sox2 bind synergistically to the Sox/Oct motif with higher affinity (Ambrosetti et al. 1997; Mistri et al. 2015), while other pluripotency genes co-occupy their targets without such known synergistic effect. Since DamID can also detect transient or weak protein-DNA interaction (Aughey and Southall 2016), DamID-specific peaks may reflect the sensitive nature of this technology. In agreement, the overlapping peaks represented the stronger peaks (i.e. peaks containing higher number of aligned reads) among peaks detected by DamID-seq (Supplemental Fig. S5).

**Supplemental Figure 5:**
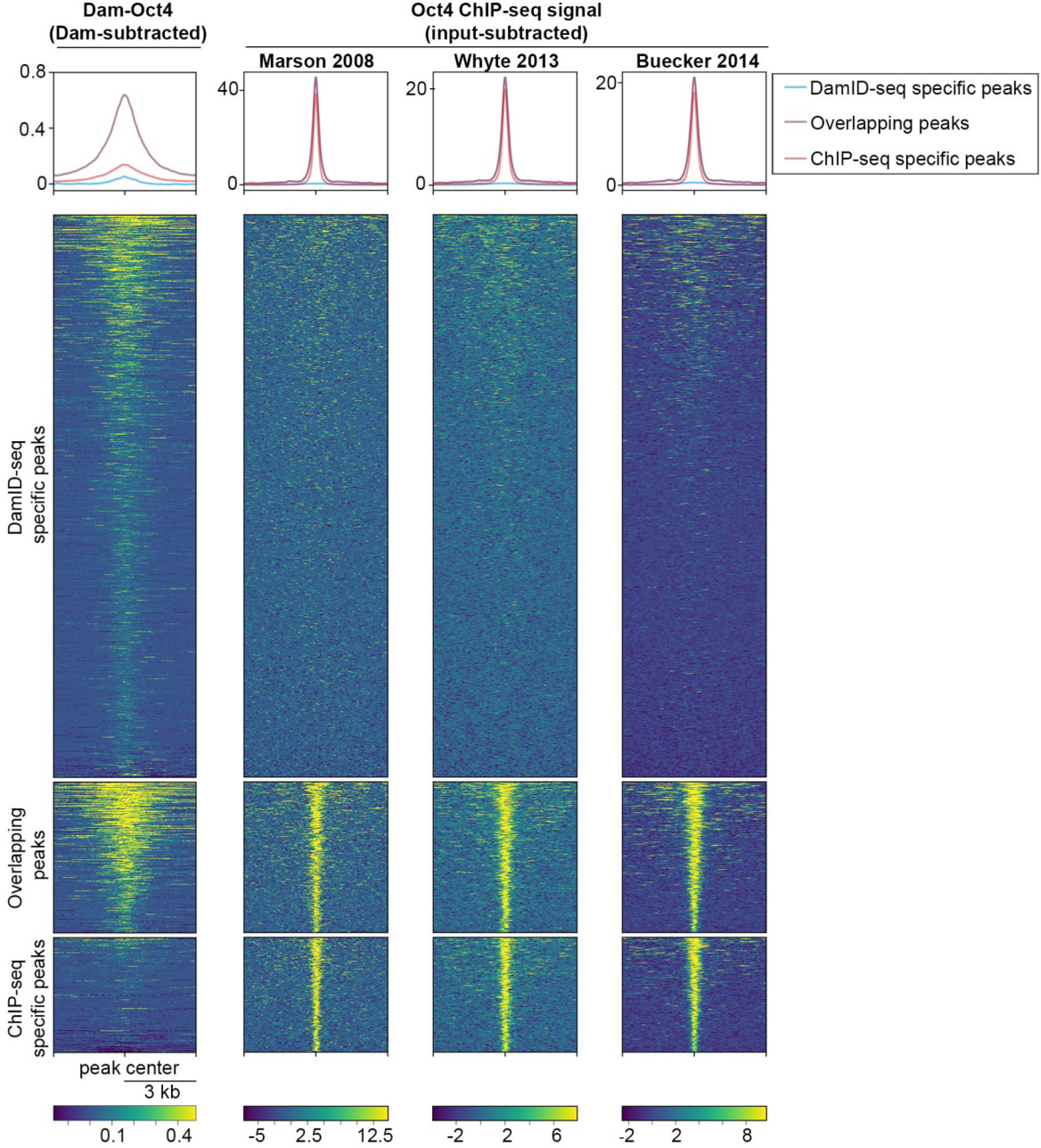
Intensity of DamID-seq and ChIP-seq signal in the overlapping and non-overlapping peaks. The heatmap shows the Dam-Oct4 methylation signal subtracted with Dam-only methylation signal in 10^6^ ESCs, and the Oct4 ChIP-seq signal (input-subtracted) from three different ChIP-seq studies (Buecker et al. 2014; Marson et al. 2008; Whyte et al. 2013). Top, middle and bottom panels represent genomic loci with Oct4 binding peaks identified by only DamID-seq, both and only ChIP-seq, respectively.Enriched gene ontology (GO) terms associated with the overlapping peaks as well as ChIP-seq specific peaks included ‘stem cell maintenance’ (Fig. 1G). Oct4 DamID-seq specific peaks were associated with genes with GO terms ‘developmental induction’ and ‘cell-cell signalling involved in cell fate commitment’ (Fig. 1G), supporting the idea that in self-renewing ESCs Oct4 transiently/weakly binds to targets important for embryo development where Oct4 also plays critical roles (Niwa et al. 2000; Simandi et al. 2016).

Enriched gene ontology (GO) terms associated with the overlapping peaks as well as ChIP-seq specific peaks included ‘stem cell maintenance’ (Fig. 1G). Oct4 DamID-seq specific peaks were associated with genes with GO terms ‘developmental induction’ and ‘cell-cell signalling involved in cell fate commitment’ (Fig. 1G), supporting the idea that in self-renewing ESCs Oct4 transiently/weakly binds to targets important for embryo development where Oct4 also plays critical roles (Niwa et al. 2000; Simandi et al. 2016).

Oct4 DamID has been recently used to validate a DamID-seq protocol in a different study (Jacinto et al. 2015). The data from Jacinto et al. presented lower signal-to-noise ratio compared to our data (Supplemental Fig. S6), probably due to the lack of DpnII digestion step before PCR amplification in their protocol and potentially reflecting difficulty in achieving optimal expression levels of Dam-only/Oct4 using viral transduction.

**Supplemental Figure 6:**
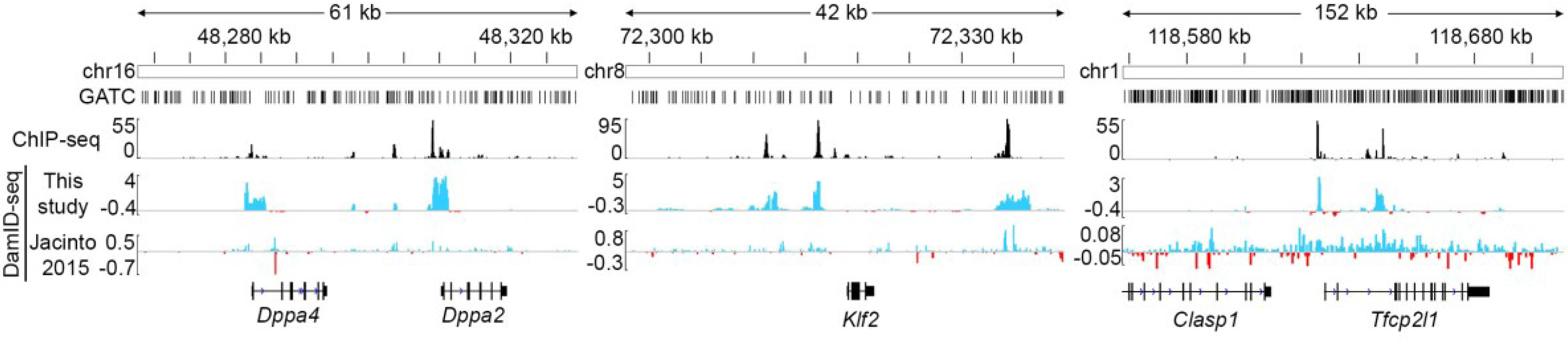
Comparison of Oct4 DamID-seq signal in different studies. Oct4 binding signal detected by ChIP-seq (Buecker et al. 2014), DamID-seq in this study with 10^6^ cells and in Jacinto et al. 2015. The protocol described in this study improves the signal-to-noise ratio of the DamID-seq signal providing high agreement with ChIP-seq.

### Transcription factor DamID-seq in 1,000 flow-sorted ESCs and NSCs

Since the DamID protocol does not include any precipitation procedure, which causes loss of DNA, we reasoned that it could be applied to lower numbers of cells. Accordingly, we performed Oct4 DamID-seq with 10^4^ and 10^3^ Rosa26-Neo-Dam and Rosa26-Neo-Dam-Oct4 ESCs (Fig. 2).

**Figure 2:**
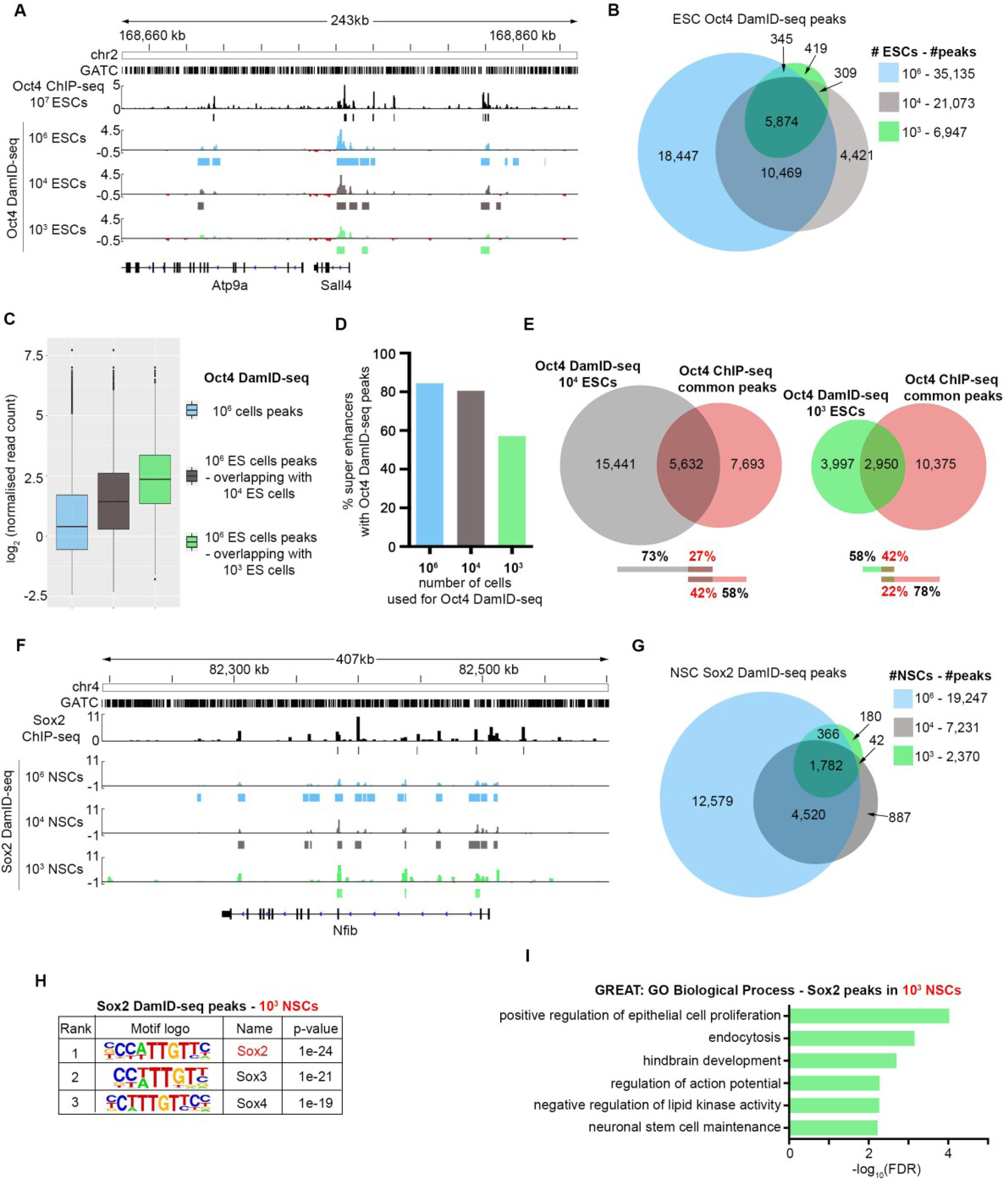
Oct4 DamID-seq in 1,000 ESCs and Sox2 DamID-seq in 1,000 NSCs. (*A*) Oct4 DamID-seq tracks from 10^6^/10^4^/10^3^ ESCs and Oct4 ChIP-seq track from 10^7^ ESCs (Buecker et al. 2014). (*B*) Overlaps of 10^6^/10^4^/10^3^ ESC Oct4 DamID-seq peaks. (*C*) Read counts of peaks in 10^6^ ESC DamID-seq (blue), and those identified by 10^4^ (gray), 10^3^ (green) ESC DamID-seq. (*D*) Percentage of ESC super enhancers (Whyte et al. 2013) containing Oct4 DamID-seq peaks using different number of cells. (*E*) Overlaps between the Oct4 ChIP-seq common peaks and DamID-seq peaks (10^4^, 10^3^ ESCs). (*F*) Sox2 DamID-seq tracks from 10^6^/10^4^/10^3^ NSCs and Sox2 ChIP-seq tracks generated from 5x106 NSCs (Mateo et al. 2015). (*G*) Overlaps of 10^6^/10^4^, or 10^3^ NSC Sox2 DamID-seq peaks. (*H*) Motif enrichment analysis (Heinz et al. 2010) of the Sox2 DamID-seq peaks from 10^3^ NSCs. (*I*) Gene ontology enrichment analysis of Sox2 DamID-seq peaks from 10^3^ NSCs using GREAT (McLean et al. 2010).

While the number of statistically significant Oct4 peaks decreases when using fewer cells, the peaks highly overlap with those identified from 10^6^ ESCs (Fig. 2A-B). Peaks identified from 10^4^ and 10^3^ cells correspond to those with higher number of reads in 10^6^ cell DamID-seq (Fig. 2C), and include ~80% and ~60% of ESC super enhancers (Whyte et al. 2013) (Fig. 2D) and 42% and 22% of ChIP-seq peaks from 10^7^ cells (Fig. 2E). To further validate this technology in different cell types and for different TFs, we performed Sox2 DamID-seq in mouse neural stem cells (NSCs) differentiated *in vitro* from Rosa26-Neo-Dam and Rosa26-Neo-Dam-Sox2 ESC lines. Sox2 binding sites identified from 10^6^, 10^4^ and 10^3^ cells highly overlapped (Fig. 2G), as seen for Oct4 DamID-seq in ESCs. The Sox2 motif was enriched in the DamID-seq peaks, and GO terms enriched in the peak-associated genes included ‘hindbrain development’ and ‘neural stem cell maintenance’ with peaks identified from 10^3^ NSCs (Fig. 2H-I). These data indicate the wide applicability of DamID-seq across different cell types and TFs, even when the starting cell number is limited. To our knowledge, these data represent the lowest number of cells used for the identification of TF binding sites in mammalian cells.

### Oct4 DamID-seq *in vivo*

Obtaining a large number of cells of interest from *in vivo* samples is often challenging, and this represents one of the major obstacles to the identification of TF targets using ChIP-seq. Our data in ESCs and NSCs suggested that DamID-seq could circumvent this limitation. Thus, we performed *in vivo* Oct4 DamID-seq using ~7.5 days post coitum (dpc) embryos containing ~15,000 cells (Snow 1977; Tzouanacou et al. 2009), where endogenous *Oct4* is ubiquitously expressed (Rosner et al. 1990; Peng et al. 2016), by generating chimeric embryos with Rosa26-Neo-Dam and Rosa26-Neo-Dam-Oct4 ESCs (Fig. 3A).

**Figure 3:**
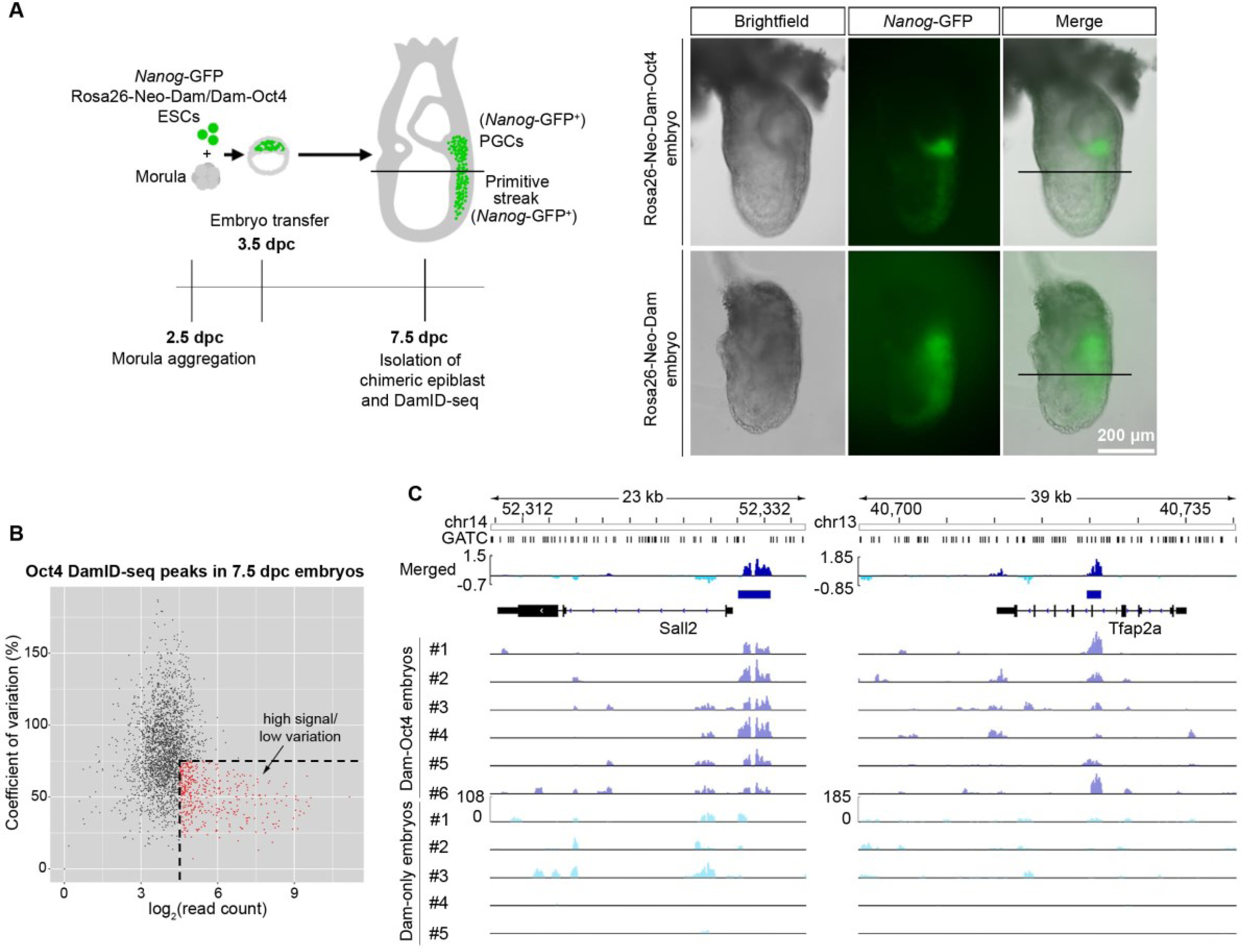
Oct4 DamID-seq with 7.5 dpc epiblasts. (*A*) DamID-seq samples were prepared from each Dam/Dam-Oct4 expressing 7.5 dpc chimeric embryo generated via morula aggregation, excluding the PGC containing region. Nanog-GFP confirms contribution of Dam/Dam-Oct4 ESCs, while the reporter expression is limited to posterior. (*B*) Read counts and coefficient of variation of the epiblast Oct4 DamID-seq peaks. Peaks with high read counts (log_2_ > 4.5) and low standard deviation (<75%) indicated in red were used for further analyses. (*C*) The merged epiblast Oct4 DamID-seq tracks (top) generated from 6 Rosa26-Neo-Dam-Oct4 and 5 Rosa26-Neo-Dam embryos. Blue bars indicate selected confident peaks from *B*.

Each embryo showed a different level of ESC contribution as observed by the amount of Nanog-GFP reporter positive cells in the posterior region of the embryos. The contribution of multiple cell types within each embryo could further reduce the consistency of Oct4 binding sites in the bulk samples. Accordingly, the in vivo DamID-seq data showed a higher variability between replicates compared to the Oct4 ESC and Sox2 NSC data (Fig. 3B, 3C, Supplemental Fig. S7). Despite this variability, we identified 2,768 statistically significant (FDR < 0.1) Oct4 DamID-seq peaks using 6 Rosa26-Neo-Dam-Oct4 and 5 Rosa26-Neo-Dam embryos as starting material, including 343 peaks with a high signal and low variability (Fig. 3B, 3C).

**Supplemental Figure 7:**
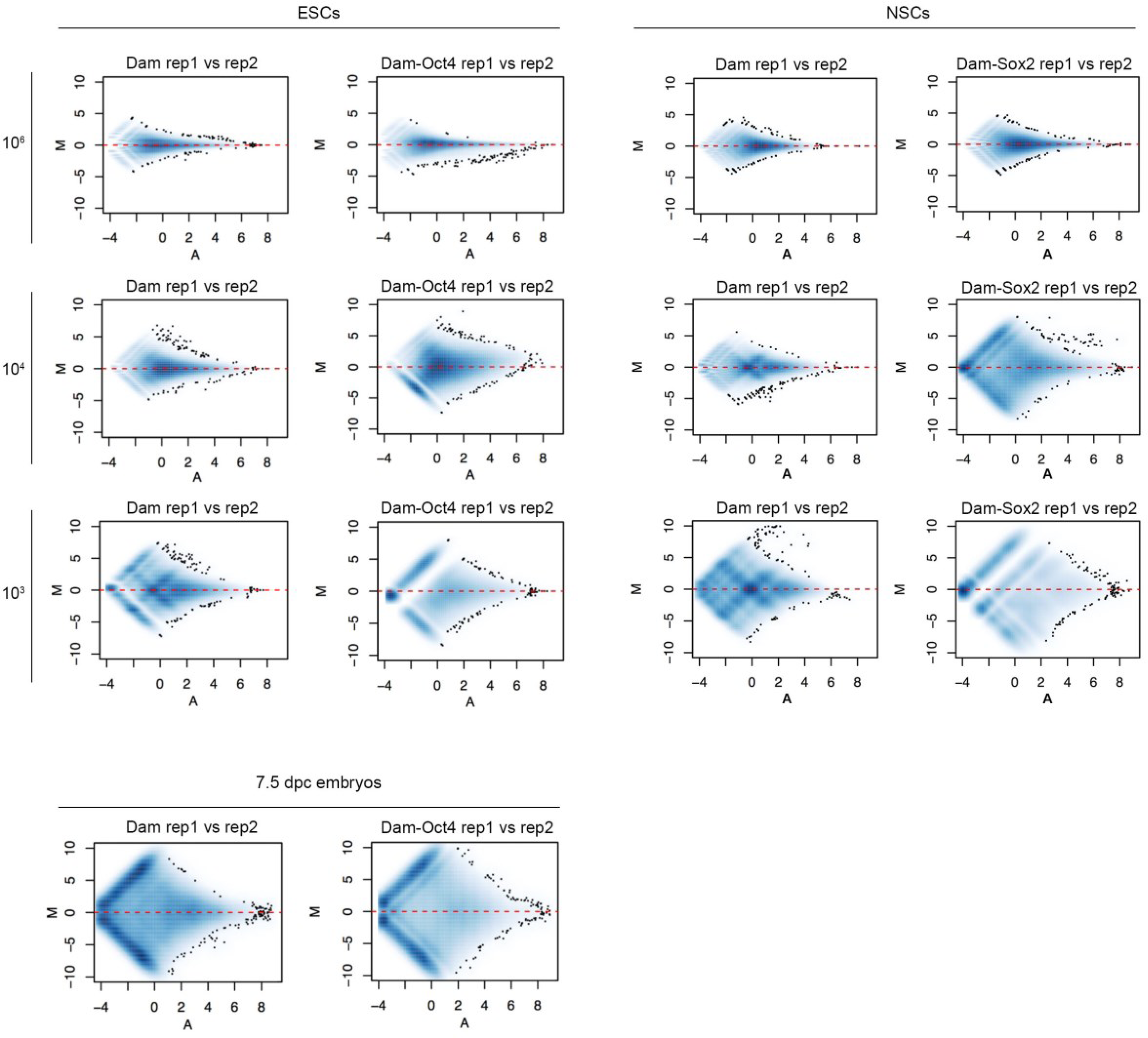
MA plots of Dam only, Dam-Oct4 and Dam-Sox2 replicates using different cell numbers and 7.5 dpc embryos. M (Y axis) represents the log_2_ of the intensity ratio of each GATC fragment between two replicates; A (X axis) represents the average log intensity of each GATC fragment. When the number of starting cells decreases, the absolute values of intensity ratio become large, indicating increasing variability between replicates.

Importantly, the Oct4 motif and GO terms related to early embryo development were highly enriched in those top 343 high-confidence Oct4 peaks and peak-associated genes, respectively (Fig. 4A, 4B). Of the 94 ‘early embryo development’ peak-associated genes, many were highly expressed in EpiSCs, the *in vitro* counterpart of the 6-7.5 dpc epiblast (Tesar et al. 2007), rather than in ESCs (Fig. 4C). Consistently, in *vivo* transcriptome data (Kojima et al. 2014) demonstrated that most of these genes were up-regulated at post-implantation stages, in particular at the early bud (EB) and late bud (LB) stages (~7.5 dpc) (green bar in Fig. 4D, Supplemental Table S2). It is noteworthy that the expression levels of the peak-associated genes, *n-Myc, Hand2, Prickle2, Sox4, Prrx1,* and *Nr2f2* changed ~2-fold 24 hours after deletion of *Oct4* at ~7.5 dpc using the Cre-loxP system (Fig. 4E, 4F) (DeVeale et al. 2013). Thus Oct4 binding signatures identified by DamID-seq provide a valuable resource to elucidate how OCT4 controls the expression of genes critical for embryo development (Osorno et al. 2012; DeVeale et al. 2013; Aires et al. 2016). Overall, we optimized a powerful technique to identify TF-DNA interaction, DamID-seq, for mammalian cells and established it as a unique strategy to identify TF binding sites in limited numbers of cells including *in vivo* samples.

**Figure 4.**
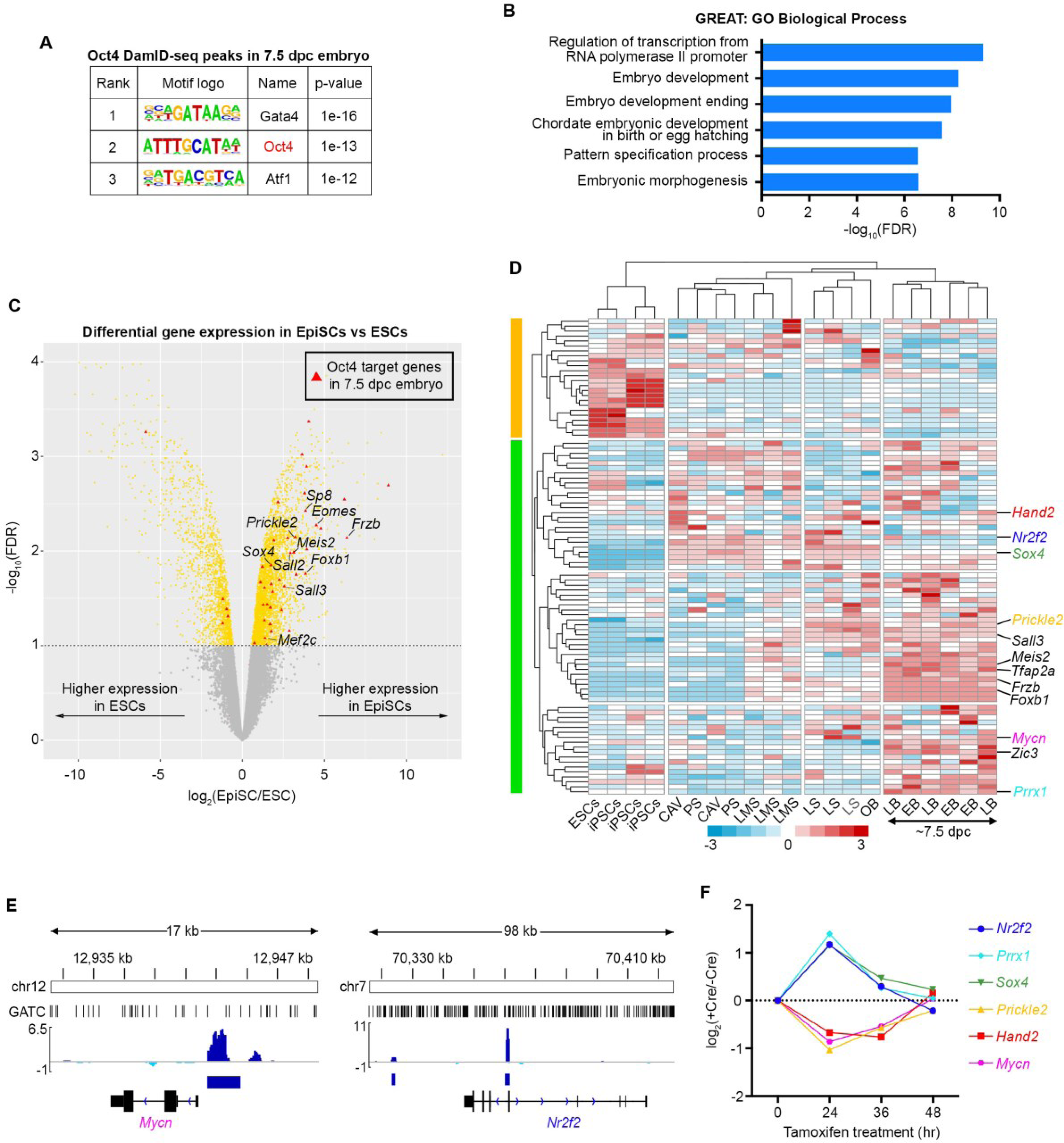
(*A*) Motif enrichment analysis (Heinz et al. 2010) of the epiblast Oct4 DamID-seq peaks. (*B*) Gene ontology enrichment analysis of Oct4 DamID-seq peaks from 7.5 dpc epiblasts using GREAT (McLean et al. 2010). (*C*) Differentially expressed genes between ESCs and EpiSCs (Tesar et al. 2007). Red triangles represent the Oct4 binding peak-associated genes in 7.5 dpc epiblasts. (*D*) Expression levels of the 7.5 dpc epiblast Oct4 binding peak-associated genes in ESCs and post-implantation epiblasts (Kojima et al. 2014). Genes whose expression significantly changed 24 hr after Oct4 deletion in the 7.5 dpc mouse embryo (DeVeale et al. 2013) are indicated in colour. CAV = epiblast of cavity; PS = prestreak; LMS = late mid streak; LS = late streak; OB = no bud; EB = early bud; LB = late-bud. (E, F) Oct4 binding peaks identified by DamID-seq (*E*) and expression changes of the development related 6 genes upon Oct4 deletion in 7.5 dpc embryos (*F*) (DeVeale et al. 2013).

## Discussion

In this work we have described an optimized DamID-seq for TF target identification in mouse cells by combining the use of the ubiquitously active endogenous *Rosa26* promoter with translation reinitiation. This approach has enabled us to detect for the first time TF targets in as few as 1,000 cells and in the developing mouse embryos at the gastrulation stage. To date ChIP-seq has been almost the only strategy used to uncover TF targets in a genome-wide manner in mammalian cells and several technical advances have been made (Furey 2012; Schmidl et al. 2015; Lara-Astiaso et al. 2014; Savic et al. 2015). Nevertheless TF ChIP-seq with less than 10,000 cells has not been reported. In *Drosophila,* DamID has been proven to be a powerful tool to investigate binding sites of, mainly but not only, chromatin proteins (van Bemmel et al. 2013). The specific binding of Dam-POI at closed chromatin stands out when compared with Dam-only expressing cells since Dam-only methylates closed chromatin less frequently (Kladde and Simpson 1994). In fact, the first successful DamID experiments in human cells were performed for heterochromatin binding protein HP1β and LaminB1 (Vogel et al. 2006; Guelen et al. 2008). More recently, DamID-seq has been used to investigate laminB1 binding even in single haploid human cells (Kind et al. 2015). In contrast, TF DamID has been more technically burdensome since Dam-only frequently methylates GATC at open chromatin loci, resulting in high background methylation signal. In addition, TFs binding sites are much narrower (8-20 bp) compared to lamina-associated domains (LADs, median size ~0.5 Mb). Therefore, the number of DNA fragments obtained from one binding domain/site following DpnI digestion is much lower. This makes TF target detection with small numbers of cells much more difficult in DamID. Thus, it is critical to express Dam-only and Dam-TF at similar expression levels, and the use of the *Rosa26* promoter together with translation reinitiation represents a successful approach. Our optimized DamID-seq has enabled the identification of TF targets using 10,000-fold less starting cells than standard ChIP-seq, and 10-to 100-fold less than the best reported TF ChIP-seq. The applicability of TF DamID-seq to much smaller cell numbers opens opportunities for new investigation in various biological contexts, even though the resolution of DamID-seq is lower than ChIP-seq as the signal relies on the frequency of the GATC sequences (~260 bp on average in the mouse genome). The PhiC31 RMCE system allows the generation of Dam-POI ESC lines *via* simple plasmid transfection. The resulting ESCs can be used to generate chimeric mice or mouse lines, making DamID-seq possible with any tissue of the mouse embryo since the Rosa26 promoter is ubiquitously active.

In addition, although the *Rosa26* promoter is constitutively active, we could successfully detect gain and loss of TF binding signatures during cell differentiation was methylation signal were diluted through cell division. When investigating TF binding dynamics in post-mitotic cells or slowly dividing cells, a possible limitation of our DamID-seq system is the lack of inducibility of Dam-only/POI protein. A Cre-loxP mediated, tissue specific Dam-only/POI protein expression system in adult mouse would further increase the applicability of DamID-seq *in vivo* (Pindyurin et al. 2016; Southall et al. 2013). About 50% of Oct4 peaks identified in 3 different ChIP-seq data sets were also identified by DamID-seq using smaller numbers of ESCs. This overlap is significant considering the intrinsic differences of the two techniques, the distinct data analysis methods, the variety of cell culture conditions and the fact that the overlap between ChIP-seq data sets themselves is also limited. In particular, stronger Oct4 DamID-seq peaks highly overlapped with the ChIP-seq peaks, as they probably represent highly robust and/or frequent binding events. Thus selection of the strongest peaks amongst the total DamID peaks may enable one to select the peaks likely detectable also in ChIP-seq. On the other hand, amongst the weaker peaks, DamID may provide insight into transient interactions that cannot be detected by ChIP-seq such as pioneering/scanning activity. In summary, DamID-seq in mammalian cells represents a novel powerful tool to reveal targets of TFs in yet unexplored biological contexts.

## Methods

### Vector construction

The coding sequence of Dam protein was taken from the pIND-V5-EcoDam (gift from Prof. Bas van Steensel, NKI). The *Neo-Dam, Neo-Dam-Oct4, Bsd-Dam, Bsd-Dam-Oct4, Hyg-Dam* and *Hyg-Dam-Oct4 pENTR* vectors were generated by Gibson assembly (Gibson et al. 2009). Two stop codons (TAA TAA) were added at the end of each antibiotic resistance genes, followed by a single C base and the ATG codon of Dam-only/Dam-Oct4 (similar to the system described by Southall, *et al.,* 2013. A 16 amino acid residue long linker (SGGGGSGGGGSGGGGS) was added between the Dam and the Oct4 coding region. *Neo-Dam* and *Neo-Dam-Oct4 pENTR* vectors were transferred into the *pROSA26-DEST* vector, after the *PGK-Neo* cassette was removed by *in vitro* Cre-loxP recombination, via Gateway LR II Clonase (Invitrogen) recombination. The *Dam\** and *Dam*-Oct4* (Dam* = D181A mutant) sequences were generated via site directed mutagenesis and transferred into a *PB-CAG-pA-DEST* (gift from Prof. Andras Nagy, Mount Sinai Hospital) vector to generate *CAG-Dam*-IRES-Puromycin-pA* and *CAG-Dam*-Oct4-IRES-Puromycin-pA.* All the plasmids and sequences are available upon request.

### Generation of cell lines via gene targeting and RMCE

The Rosa26-Neo-Dam, Rosa26-Neo-Dam-Oct4, Rosa26-BSD-Dam and Rosa26-Bsd-Dam-Oct4 ESC lines were generated via standard gene targeting strategy; the *pROSA26-ATTP-Neo-Dam-ATTP, pROSA26-ATTP-Neo-Dam-Oct4-ATTP, pROSA26-ATTP-Bsd-Dam-ATTP, pROSA26-ATTP-Bsd-Dam-Oct4-ATTP* vectors generated with *pRosa-DEST* (Addgene #21189) were used for the homologous recombination in the *Rosa26* locus of the TNG-MKOS cell line (Chantzoura et al. 2015). The clones with the successful targeting event were identified via southern blot using HindIII-or PacI-digested genomic DNA for the Rosa26 probe or the Neo/Bsd probe, respectively. The following primers were used to generate the three probes:

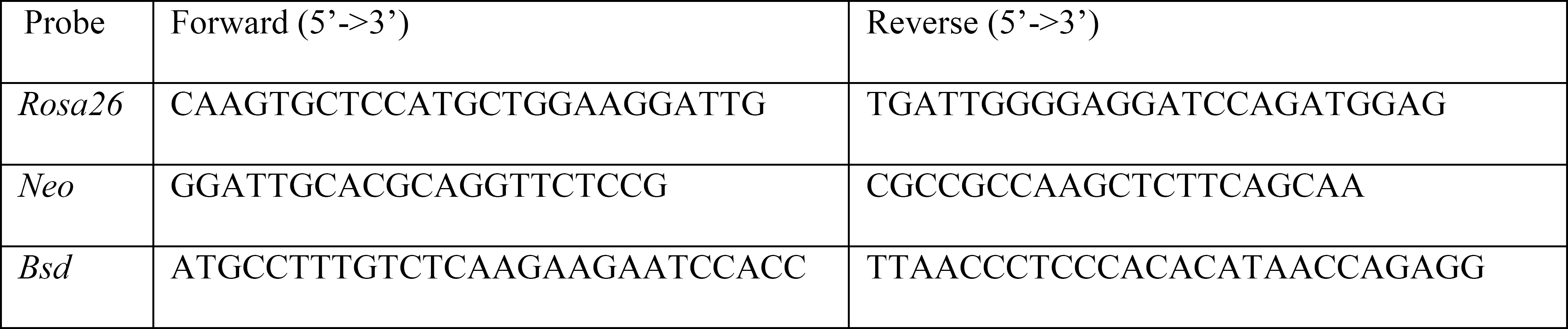

For the φC31-mediated cassette exchange, Rosa26-Bsd-Dam-Oct4 (4x10^5^) cells were plated on a gelatinized well of a 6-well dish. 24 hours later, a total of 7 μg of DNA (3.5 µg of vector encoding for φC31 integrase (gift from Prof. Andras Nagy, Mount Sinai Hospital) and 3.5 μg of *ATTB-Hyg-Dam-ATTB-pENTR* or *ATTB-Hyg-Dam-Oct4-ATTB-pENTR*) was used for each transfection using Lipofectamine3000 (Invitrogen). 24 hours after lipofection, cells were replated onto 5 gelatinized 55 cm^2^ dishes and hygromycin (Hyg) selection (150 μg/ml) was started on the next day. Hyg resistant colonies were picked 10 days later, and the successful recombination event was confirmed by genomic PCR using the following primers.

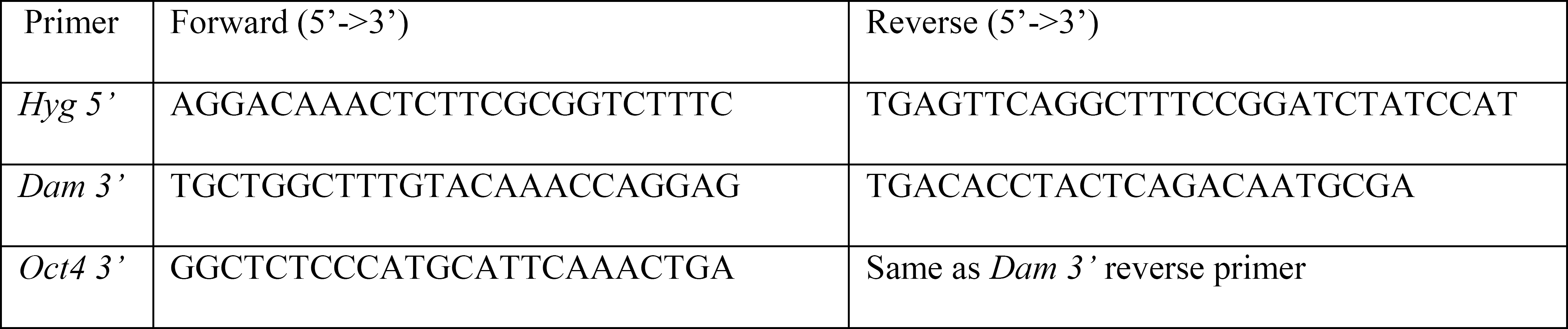

### ESC culture

ES cells were in Glasgow minimum essential medium (GMEM) supplemented with 10% fetal calf serum (FCS), penicillin-streptomycin, 1x non-essential amino acids (Invitrogen), 1 mM sodium pyruvate, 2 mM glutamine, 0.05 mM β-mercaptoethanol (Life Technologies) supplemented with human leukemia inhibitory factor (LIF), 100 U/ml on gelatinized flasks in a 37 °C/5% CO_2_ incubator. For the 2i condition, cells were grown in N2B27 supplemented with µM PD0325901 and 3 µM CHIR99021 (Axon Medchem).

### ESCs to EpiLCs differentiation

Rosa26-Neo-Dam and Rosa26-Neo-Dam-Oct4 ESCs were cultured in serum-free N2B27-based medium (containing DMEM/F12 medium supplemented with N2 combined 1:1 with Neurobasal® medium supplemented with B27; all from Thermo Fisher Scientific), MEK inhibitor (PD0325901, 0.8 μM, Axon Medchem) and GSK3b inhibitor (CHIR99021, 3.3 μM, Axon Medchem) on gelatinized tissue culture dishes (Ying et al. 2008) for at least 5 passages. For EpiLC differentiation, 100,000 cells were replated onto fibronectin-coated (7.5 μg/ml) 22 mm dish in medium serum-free N2B27-based medium supplemented with 1% KOSR, 2 μM glutamine (Invitrogen), 0.05 mM β-mercaptoethanol (Thermo Fisher Scientific), 1x non-essential amino acids (Invitrogen), 10 ng/ml FGF2 (R&D), 20 ng/ml ActivinA (Peprotech) for 72 hours before harvesting.

### ESCs to NSCs differentiation

NSC lines were derived from ESCs as previously described (Pollard et al. 2006). Specifically, *Rosa26-Neo-Dam* and *Rosa26-Neo-Dam-Sox2* ESCs maintained in 2i medium were initially plated on gelatin-coated 6-well plates at 10^6^, 7.5x10^5^ and 5 x10^5^/well in NSC complete medium containing DMEM/F-12 Media, 1:1 Nutrient Mixture (Sigma, D8437), 1X N2 supplement (Thermo Fisher Scientific), 1X B27 supplement (Thermo Fisher Scientific), 8 mM glucose (Sigma, G8644), penicillin-streptomycin, 0.001% BSA (ThermoScientific, 15260037), 0.05 mM β-mercaptoethanol (Thermo Fisher Scientific). Cells were fed daily for 7 days. On day 7, the wells that have more cells with NSC-like morphology were selected, and 1/2 of the cells were seeded into a non-gelatinised flask in NSC complete medium supplemented with 10 ng/ml mouse EGF (Peprotech) and 10 ng/ml human FGF2 (Peprotech). Cells were left to form floating aggregates for 3 days. On day 10, aggregates were harvested by centrifugation of the supernatant at 700 rpm for 3 minutes. Cells were then resuspended in NSC complete media supplemented with EGF, FGF2 and 1 μg/ml Laminin (L2020, Sigma) and seeded onto non-gelatinized T25 flasks. Aggregates were allowed to settle over the course of 10 days, with medium changed every 2-3 days. On day 20, when the flask reached ~50% confluence, half of the cells were passaged into T25 flasks and defined as p1 NSCs. NSCs were maintained in EGF, FGF2, Laminin-supplemented NSC complete medium and passaged every 3-4 days at 1:6/1:8 density. To ensure homogeneity, cells were passaged at least 4 times before performing DamID-seq.

### Dissection of 7.5 dpc epiblasts

Rosa26-Neo-Dam and Rosa26-Neo-Dam-Oct4 chimeric embryos generated by morula aggregation were collected at 7.5 dpc. After removing the Reichert’s membrane, embryos were imaged to evaluate the presence of *Nanog*-GFP positive cells. The distal part of epiblast was dissected with sharp glass needles in M2 (Sigma, M5910) medium to exclude primordial germ cells. Each epiblast was washed once with PBS, transferred into the Genomic Lysis Buffer provided in the Quick-gDNA™ MicroPrep and processed separately as described in the DamID-seq section for the generation of DamID-seq libraries.

### *Oct4^-/-^* ESC rescue experiment and Alkaline Phosphatase (AP) staining

Rescue plasmids with *Oct4, Dam*-Oct4, Dam\** cDNA followed by the *IRES-Puromycin* resistance gene cassette under the CAG promoter were transfected into ZHBTc4 cells (Niwa et al. 2002) using Lipofectamin 3000 (Life Technologies). One day after transfection, cells were replated and cultured in the presence of puromycin (1 μg/ml) and doxycyxline (1 μg/ml) for 12 days. Then, the cells were fixed, washed in distilled water and stained for alkaline phosphatase (AP) expression using a leukocyte alkaline phosphatase kit (Sigma).

### ESCs to fibroblast differentiation

Fifty thousand Rosa26-Neo-Dam and Rosa26-Neo-Dam-Oct4 ESCs were plated in a gelatin-coated 35 mm dish in ESC medium without LIF with 1 μM retinoic acid (Sigma) and cultured for 9 days. RNA was extracted from the *in vitro* differentiated fibroblasts and used for qRT-PCR for the expression analysis of mRNA coding Dam protein.

### RNA isolation, cDNA synthesis and quantitative real-time RT-PCR

RNA was isolated using TRIzol (Life Technologies) according to manufacturer’s instructions. The RNA was then reverse transcribed to cDNA with the Moloney Murine Leukemia Virus Reverse Transcriptase (M-MLV RT, Life Technologies). The qPCR reactions were performed in 384 well plates, in a 10 μl reaction containing cDNA corresponds to 10 ng of starting RNA, 0.5 μM primers and 5 μl LightCycler® 480 SYBR Green I Master Mix (Roche) using LightCycler® 480 (Roche). The primers used for gene expression analyses are:

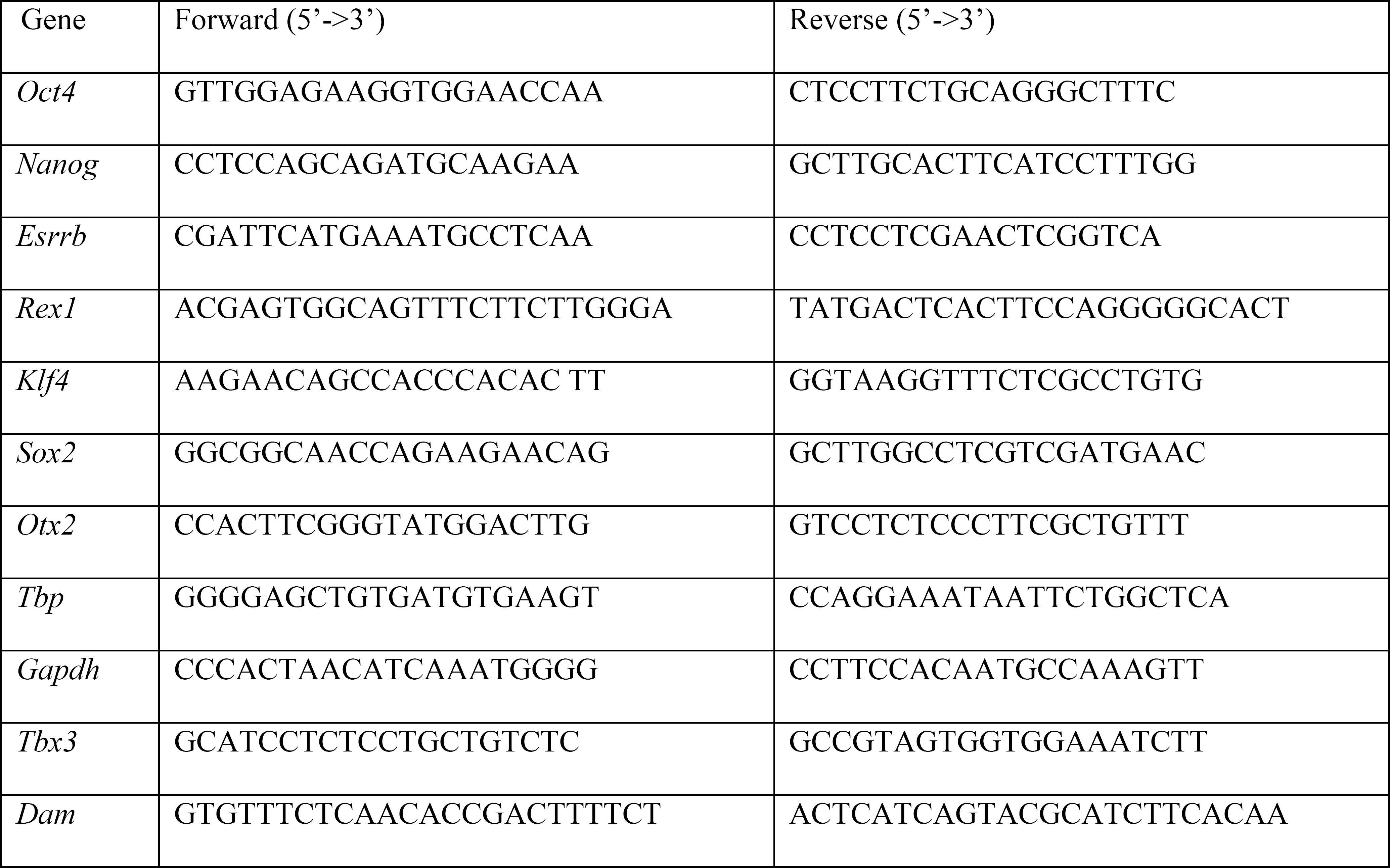

### Quantitative DamID (qDamID)

Dam-only/Dam-Oct4 protein expressing cells (~1-2x10^6^) were collected via trypsinization and genomic DNA (gDNA) was extracted using the DNeasy Blood & Tissue Kit (Qiagen) according to the manufacturer’s instructions. Two microgram of gDNA were diluted in 16 μl of H_2_O. The gDNA solution was then splitted in two tubes (8 μl of DNA solution in each tube): in one tube the DpnII buffer (1 μl, 1X final) and the DpnII enzyme (1 μl, NEB) were added, in the other tube only the DpnII buffer (1 μl, 1X final) and H_2_O (1 μl) were added. Both tubes were incubated overnight at 37 °C. gDNA was then diluted fifty times to get a final concentration of 2 ng/μl and 5 μl (10 ng of digested gDNA in total) were used in the qPCR analysis in three technical replicates. The qPCR reactions were performed in 384 well plates, in a 10 μl reaction including 4.5 μl of LightCycler® 480 SYBR Green I Master Mix, 0.5 μl of forward primer (10 μM) and 0.5 μl of reverse primer (10 μM), using the LightCycler480 instrument (Roche). The qPCR signal was converted to an absolute numerical value based on an standard curve, and the average of the triplicates from digested (+DpnII) and undigested (-DpnII) gDNA was used to calculate the percentage (%) of methylation at GATC_x_ site in Dam-only or Dam-Oct4 expressing cells:

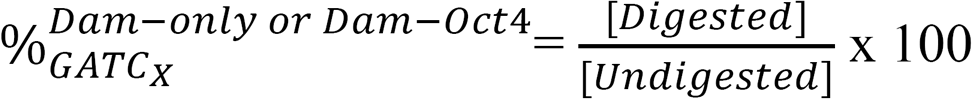

This value was the used to estimate the enrichment of Oct4 over Dam 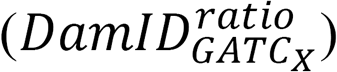 expressed as a log_2_ ratio:

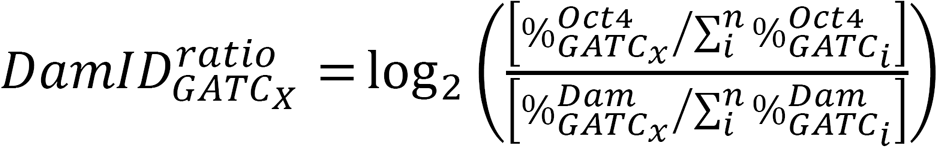

The primers used for the qDamID are listed in the table below.

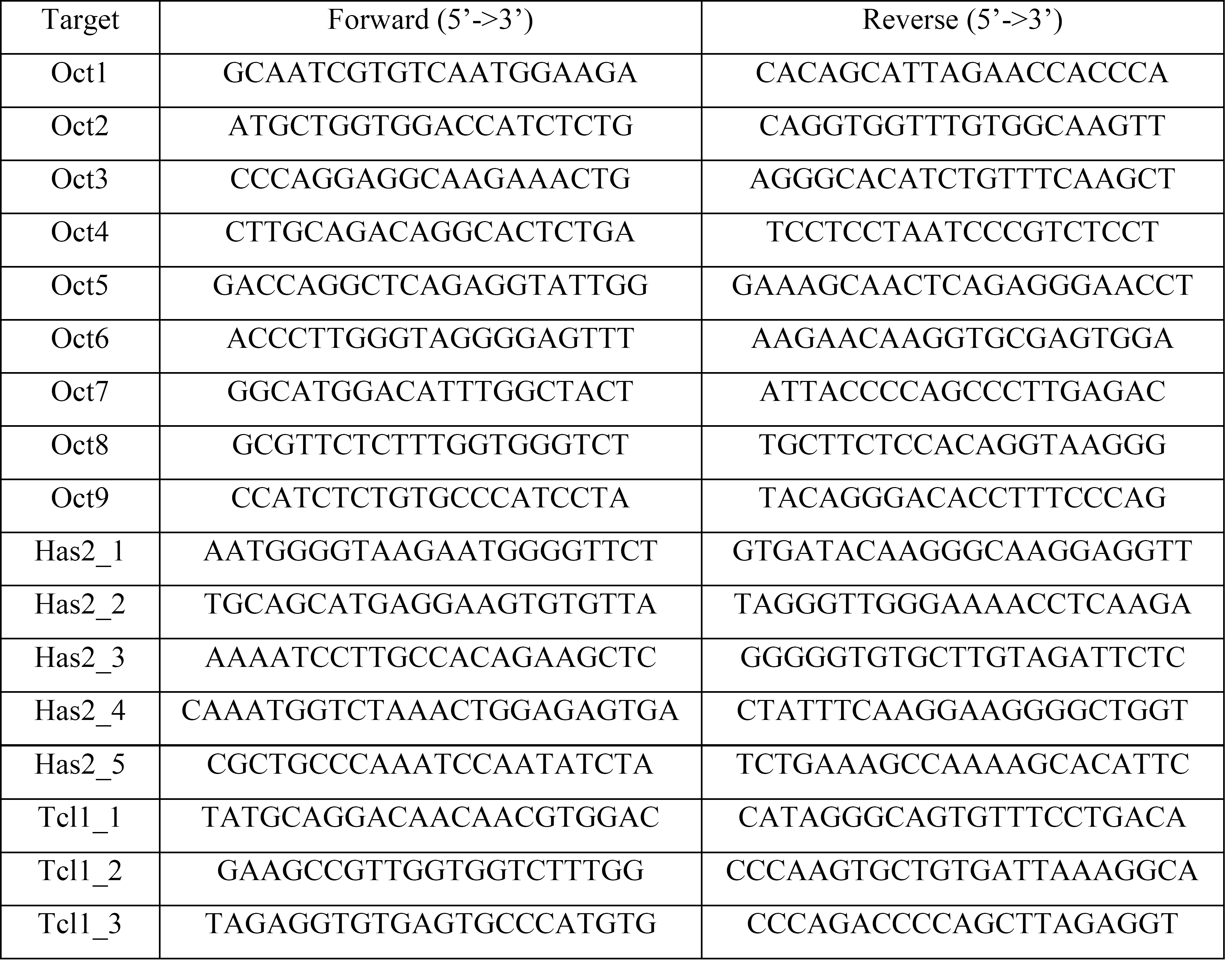

### DamID-seq

The desired number of cells where FAC-sorted using BD FACSAriaII. When the number of cells was above 10,000, cells were first spun down and then resuspended in the Genomic Lysis Buffer provided in the Quick-gDNA™ MicroPrep (ZymoResearch). When number of cells was lower than 10,000, cells were sorted directly into the Genomic Lysis Buffer. The gDNA was extracted according to the manufacturer’s instructions and eluted with 10 μl of the Elution Buffer. The gDNA was transferred into a 0.2 ml PCR tube containing 5 μl of DpnI mix (20 units DpnI, 1x Cutsmart buffer [NEB]), and digested at 37 °C for three hours before heat inactivation of the enzyme at 80 °C for 20 minutes. Double-strand DamID adapters (Vogel et al. 2007) were prepared by annealing AdRt (5’-CTAATACGACTCACTATAGGGCAGCGTGGTCGCGGCCGAGGA-3’, IDT) and AdRb (5’-TCCTCGGCCG -3’, IDT). Adapter ligation was performed by adding 5 μl of ligation mix (20 units T4 Ligase, 1x T4 Ligase Buffer [NEB] and 0.2 µM DamID adapters) to the 15 μl of DpnI digested DNA solution and keeping at 16 °C overnight. After heat inactivation of ligase at 65 °C for 20 minutes, 5 μl of DpnII mix (10 unit of DpnII and 1x DpnII buffe [NEB]) were added and the samples were kept at 37 °C for one hour before heat inactivation at 65 °C for 20 minutes. Then, 100 μl of PCR mix (KAPA HiFi HS ReadyMix [KAPA Biosystems, 1X], 10 μM AdR PCR primer, 5’-GGTCGCGGCCGAGGATC-3’ [IDT], 1X SYBR® Green I Nucleic Acid Gel Stain [Life Technologies]) were added to the tube. From the final volume of 125 μl, 10 μl were used to perform qPCR in technical duplicate (spending total 20 μl), and the number of PCR cycles to stop the PCR in the log-linear amplification phase were determined. The remaining 105 μl were then used to amplify the adapter-ligated fragments using the following program:

| Temperature (°C) | Time (min) | Cycles |
| --- | --- | --- |
| 72 | 10 | 1 |
| 94 | 1 | 1 |
| 65 | 5 |  |
| 72 | 15 |  |
| 94 | 1 | 4 |
| 65 | 1 |  |
| 72 | 10 |  |
| 94 | 1 | (Determined by qPCR for each sample) |
| 65 | 1 |  |
| 72 | 2 |  |

At the end of the PCR amplification, the samples were purified using SPRI magnetic beads. Briefly, 126 μl (1.2X DNA volume) of beads were mixed with the amplified DNA and kept for 5 minutes at room temperature before moving them onto a magnetic rack. After 5 minutes on the magnetic rack, the supernatant was removed and the beads were washed twice with 80% ethanol (150 μl, 30 seconds each). After 5 minutes of open-air drying, beads were removed from the magnetic rack, resuspended in 32.5 μl H_2_O and kept at room temperature for 3 minutes. Tubes were put back on the magnetic rack for 2 minutes, and the clear supernatant (30 μl, containing the DNA of interest) was transferred into new 1.5 ml eppendorf tubes and quantified by Qubit 2.0 (Invitrogen). Agarose gel electrophoresis usually shows a smear in the range 200-2000bp. The sequencing libraries presented in this study have been prepared using the Nextera DNA® sample preparation kit (Illumina), according to the manufacturer’s instructions. Briefly, after Qubit quantification, 50 ng of the PCR amplified DNA was used in a tagmentation reaction (5 minutes) in which the Tn5 enzyme fragments the DNA and simultaneously inserts the pre-loaded adapters. After a DNA purification step using the DNA Clean & Concentrator™ Kit (Zymo Research), the tagmented DNA was amplified by PCR for 5 cycles using the following program:

| Temperature (°C) | Time | Cycles |
| --- | --- | --- |
| 72 | 3 min | 1 |
| 98 | 10 sec | 5 |
| 63 | 30 sec |  |
| 72 | 3min |  |

Primers used in the PCR amplification are barcoded, so a different pair of primers is used for each sample. After the PCR, DNA is purified using SPRI magnetic beads (0.8X DNA volume) and eluted in 30 μl of water. The purified DNA was used for Qubit 2.0 quantification and Tapestation analysis before pooling the libraries and submitting to the sequencing facility. The concentration of the library pool was usually ~25-50 nM; 50 bp single-end sequencing was performed at Edinburgh Genomics on the Illumina HiSeq 2,500 (chemistry v. 4) System or at BGI on the Illumina HiSeq 4,000 System. A list of all the reagents and volumes required for each step of the newly optimized DamID-seq protocol (from gDNA extraction to next-generation library preparation) is summarized in the table below.

| Step | Reagent used, company | Mix volume | Total volume of sample |
| --- | --- | --- | --- |
| **gDNA extraction** | gDNA<sup>TM</sup> MicroPrep (Zymo Research) | 10 µl | 10 µl |
| **DpnI digestion** | DpnI (NEB) | 5 µl | 15 µl |
| **Adapter Ligation** | T4 Ligase (NEB)<br>DamID Adapters (IDT) | 5 µl | 20 µl |
| **DpnII digestion** | DpnII, NEB | 5 µl | 25 µl |
| **PCR amplification** | Kapa HiFi HS RM (Kapa Biosystems)<br>DamID PCR primers (IDT)<br>SYBR Green (Life Technologies) | 100 µl | 125 µl<br>(10 µl x2 for qPCR, 105 µl for the sample amplification) |
|  |  | **DNA amount** | **Total volume of sample** |
| **Library preparation** | Nextera DNA Library Prep Kit (Illumina) | 50 ng | 30 µl |

### DamID-seq data analysis

#### Accession numbers

All the raw and processed DamID-seq data generated in this study can be accessed at the following private link:

#### Data quality, alignment, and read counting

Read quality was checked using FastQC (Andrews 2014) (version 0.11.5) and adapters were trimmed using cutadapt (Martin 2011) (version 1.13). Reads were aligned to the UCSC mm10 assembly of the mouse genome (Church et al. 2009) using bwa mem (Li and Durbin 2010) (version 0.7.15). Unmapped, secondary, and supplementary alignments were filtered using SAMtools (Li et al. 2009) (version 1.4). Reads mapped to ENCODE blacklist regions (ENCODE Project Consortium 2012), alternate, unplaced, and mitochondrial contigs were filtered using BEDtools (Quinlan and Hall 2010) (version 2.26.0). Reads were counted into fragments using the featureCounts command from Subread (Liao et al. 2013) (version 1.5.0). Fragments were defined as regions of the genome enclosed by a GATC sequence. Only reads mapped fully within a fragment were counted.

#### Count filtering, normalisation, and peak calling

The following analysis was carried out using the statistical programming language R (R Core Team 2013) (version 3.3.2) and packages from the Bioconductor project (Huber et al. 2015) (version 3.4). Fragments with 10 read counts or lower in equivalent average log count-per-million (aveLogCPM) values were removed. Normalisation factors were calculated with the normOffsets function from the csaw package (version 1.8.1). The type argument was set to “scaling”. Using these normalisation factors, log count-per-million (logCPM) values were calculated and smooth quantile normalised using the qsmooth package (version 0.0.1) The quantro package (version 1.8.0) was used to assess the appropriateness of using smooth quantile normalisation over regular quantile normalisation. Differentially bound fragments were identified by testing for differential abundance between the Dam-POI (Protein-of-Interest) and Dam-only samples. The normOffsets/qsmooth normalised count matrix was used with limma-trend from the limma package (3.30.13). The trend and robust arguments were set to TRUE. Fragments were then combined into differentially-bound regions using the mergeWindows and combineTests function from the csaw package. The tol and max.width arguments were set to 260 (median GATC fragment size for the mm10 assembly) and 10,000, respectively. Peaks were defined as regions with a false discovery rate (FDR) smaller than 0.1 and a log fold-change (logFC) greater than 0.5.

#### Visualisation

Read coverage across the genome was calculated with the genomecov command from BEDtools (version 2.26.0). Coverage for each sample was scaled using the normalisation factors calculated previously. The bedGraph files were converted into bigWig files with the UCSC bedGraphToBigWig (Kent et al. 2010) tool (version 4). Subtracted bigWig files were generated using the diff command from WiggleTools (Zerbino et al. 2014) (version 1.2).

### Downstream analysis of DamID-seq data

The intersection between different sets of peaks was calculated using the tool mergePeaks of the Homer suite (Heinz et al. 2010) (version 4.8.3). Proportional Venn diagram of three sets was generated using the software eulerApe (Micallef and Rodgers 2014) (version 3) while proportional Venn diagrams of two sets were generated using the Venn diagram generator (http://jura.wi.mit.edu/bioc/tools/venn.php). Box-plots and scatter plots were generated using the package ggplot2 in the R statistical environment (version 3.3.2). The deepTools suite (Ramírez et al. 2016) was used to generate the heatmap in Figure Suppl. 5 using the computeMatrix and plotHeatmap commands.

### DNase-seq

#### Accession numbers

DNase-seq hypersensitivity data for E14 mouse embryonic stem cells were downloaded from GEO with the BioSample accession number GSM1014154.

#### Visualisation

Read coverage was lifted over from the UCSC mm9 to mm10 assembly of the mouse genome (Church et al. 2009) using CrossMap (Zhao et al. 2014) (version 0.2.2).

### ChIP-seq data set analysis

#### Accession numbers

ChIP-seq Oct4 data for E14 mouse embryonic stem cells were downloaded from GEO with the GEO DataSet accession numbers GSE11724, GSE44288 and GSE56138; ChIP-seq Sox2 data for proliferating neural stem cells were downloaded from European Nucleotide Archive, accession numbers ERR216111, ERR414096 and ERR414097.

#### Data quality, alignment, and filtering

Read quality was checked using FastQC (Andrews 2014) (version 0.11.5) and adapters were trimmed using Trim Galore! (Kruger 2012) (version 0.4.3). Multiple quality reports were aggregated and visualised using MultiQC (Ewels et al. 2016) (version 0.9). Reads were aligned to the UCSC mm10 assembly of the mouse genome (Church et al. 2009) using bwa mem (Li and Durbin 2010) (version 0.7.15). Duplicates were removed using the MarkDuplicates command from Picard (Broad Institute) (version 2.9.0). Unmapped, secondary, and supplementary alignments were filtered using SAMtools (Li et al. 2009) (version 1.4). Reads mapped to ENCODE blacklist regions (ENCODE Project Consortium 2012), alternate, unplaced, and mitochondrial contigs were filtered using BEDtools (Quinlan and Hall 2010) (version 2.26.0).

#### Peak calling and visualisation

For studies without biological replicates (GSE11724 and GSE44288) peak calling was performed with the following method: Fragment size was estimated using cross-correlation analysis from the phantompeakqualtools software package (Landt et al. 2012). Peaks were called using the callpeak command from MACS2 (Liu 2014) (version 2.1.1). The following arguments were used: -g mm –keep-dup all -s <read length> –extsize <estimated fragment size> -q 0.05 -B –SPMR.

For studies with biological replicates (GSE56138 and Sox2 ChIP-seq), peak calling was performed according to the ENCODE (Phase-3) ChIP-seq pipeline (Landt et al. 2012). Briefly, self-pseudoreplicates, pooled replicates, and pooled self-pseudoreplicates were generated for each sample. Fragment size was estimated using cross-correlation analysis from the phantompeakqualtools software package (Landt et al. 2012). Peaks were called on the self-pseudoreplicates, pooled replicates, and pooled self-pseudoreplicates using the callpeak command from MACS2 (Liu 2014) (version 2.1.1). The following arguments were used: -g mm –keep-dup all -s <read length> –nomodel –shift 0 –extsize <fragment size> -p 0.05 -B –SPMR. High confidence/reproducible peaks were identified using an IDR (Li et al. 2011) threshold of 0.05 (version 2.0.3). Peaks identified from the pooled pseudo-replicates dataset were used for the final set of peak calls.

Input-subtracted read coverage was calculated using the bigwigCompare command from deepTools (Ramírez et al. 2016) (version 2.4.3). The bigWig files were generated from the treatment pileup and control lambda bedGraph files from MACS2 using the UCSC bedGraphToBigWig tool (Kent et al. 2010) (version 4).

### Motif enrichment (Homer) and Gene Ontology (GREAT) analyses

The motif enrichment analysis was performed with the findMotifsGenome tool of the Homer (Heinz et al. 2010) suite version 4.8.3. For the motif enrichment, peaks were ranked by signal intensity and the highest 25%, (except 7.5 dpc embryo for which all the 343 peaks), were used as foreground. For the DamID-seq data and DamID-seq/ChIP-seq overlapped peaks, all Dam-only/Dam-Oct4 differentially bound regions (Dam-only enriched, Dam-Oct4 enriched including the foreground) were used as background. For the ChIP-seq data, the background was generated by the Homer software. GREAT analyses (McLean et al. 2010) were performed using GREAT v3.0 using the default “single nearest gene (within 1000kb)” association rule. The highest 25% of each set (ranked by signal intensity) was used as input for each sample; all the 343 peaks identified in the 7.5 dpc embryos were used as input for the GREAT analysis.

### Microarray analysis

The microarray dataset analysis (GEO accession number GSE46227) was carried out using the statistical programming language R (R Core Team 2013) (version 3.3.2) and packages from the Bioconductor project (Huber et al. 2015) (version 3.4). Data were normalized and log_2_-transformed by using the limma (Smyth 2004) package (version 3.28.21). The heatmap was generated by using the pheatmap package (version 1.0.8).

### Data access

All DamID-seq data have been deposited in the Gene Expression Omnibus under the accession code GSE98092.

## Acknowledgements

We thank F. Rossi and C. Cryer for assistance with flow cytometry. L. Robertson and J. Rennie for technical assistance with morula aggregations, and Biomed unit staff for mouse husbandry; F. Wymeersch for the help with the isolation and imaging of the embryos; S. Picelli for suggestions about tagmentation and G. Pavesi for the help with motif enrichment analysis. We thank A. Soufi, K. Ottersbach, S. Schoenfelder, M. Beniazza, D. F. Kaemena, U. Gunesdogan and A.C. McGarvey for comments on the manuscript. This work was supported by European Research Council (ERC) grants ROADTOIPS and MRC senior non-clinical fellowship funded for K.K. L.T. and J.A. are funded by CMVM scholarships and Principal’s Career Development scholarship from the University of Edinburgh. B.C. is supported by BBSRC EASTBIO Doctoral Training Partnership.

## Author contributions

L.T. generated the vectors and cell lines used in the study, designed and performed the qDamID/DamID-seq experiments, analyzed the data, and wrote the manuscript with input from the coauthors. J.A. performed all the analyses for the identification of DamID-seq peaks and reanalyzed the published ChIP-seq data; N.B. generated the *Rosa26-Neo-Dam-Sox2* cell line, helped with embryo collection and preparation of DamID-seq libraries. B.C. derived the NSC lines. V.W. dissected and staged the embryos. S.R.T. helped with the analysis of DamID-seq data. K.K. conceived the study, supervised experiment design and data interpretation, and wrote the manuscript.

